# GLUT1 ablation in astrocytes paradoxically improves central and peripheral glucose metabolism via enhanced insulin-stimulated ATP release

**DOI:** 10.1101/2022.10.06.511112

**Authors:** Carlos G. Ardanaz, Aida de la Cruz, Marcos Elizalde-Horcada, Elena Puerta, María J. Ramírez, Jorge E. Ortega, Ainhoa Urbiola, Cristina Ederra, Mikel Ariz, Carlos Ortiz-de-Solórzano, Joaquín Fernández- Irigoyen, Enrique Santamaría, Gerard Karsenty, Jens C. Brüning, Maite Solas

## Abstract

Astrocytes are considered an essential source of blood-borne glucose or its metabolites to neurons. Nonetheless, the necessity of the main astrocyte glucose transporter, i.e. GLUT1, for brain glucose metabolism has not been defined. Unexpectedly, we found that brain glucose metabolism was paradoxically augmented in mice with astrocytic GLUT1 ablation (GLUT1^1′GFAP^ mice). These mice also exhibited improved peripheral glucose metabolism especially in obesity, rendering them metabolically healthier. Importantly, GLUT1^1′GFAP^ mice did not present cognitive alterations. Mechanistically, we observed that GLUT1-ablated astrocytes exhibited increased insulin receptor-dependent ATP release, and both astrocyte insulin signalling and brain purinergic signalling are essential for improved brain function and systemic glucose metabolism. Collectively, we demonstrate that astrocytic GLUT1 is central to the regulation of brain energetics, yet its ablation triggers a reprogramming of brain metabolism sufficient to sustain energy requirements, peripheral glucose homeostasis and cognitive function.

## Main

The precise coupling between energy production and energy demand is still an important conundrum regarding brain metabolism. The brain is one of the most energy-demanding organs in the body and shows an urgent reliance on glucose. Indeed, despite representing only 2% of the body weight, the human brain accounts for approximately 25% of the total resting body glucose consumption^1, 2^. Arterial blood-borne glucose is the obligatory and major energy source for the brain^3^. Alternative substrates can also be metabolized by the brain, but none of them can fully replace glucose^4^. Importantly, brain glucose supply is critical for memory acquisition and consolidation at the hippocampal level^5, 6^. Besides, CNS glucose availability is also sensed by specialized hypothalamic circuits that monitor the body energy state and regulate systemic glucose metabolism accordingly^7, 8^. This suggests that central energy metabolism is intimately linked not only to local brain activity but also to peripheral metabolism.

Brain glucose supply is controlled by glucose transporters (GLUTs). Specifically, to reach brain cells, blood glucose is transported across endothelial membranes via glucose transporter 1 (GLUT1). Endothelial cells are ensheathed by the endfeet of astrocytes, whose major glucose transporter is also GLUT1^9, 10^. Thus, astrocytes are located in a privileged position to control the access of glucose into the brain^11^. Although glucose can be delivered directly to neurons, which express GLUT3 and GLUT4^12, 13^, one of the most popular theories explaining brain glucose dynamics is the astrocyte-neuron lactate shuttle (ANLS)^14^. The ANLS model postulates that neuronal activity increases astrocytic glucose uptake and glycolysis, producing lactate. This lactate is subsequently exported to the extracellular space and imported by neurons via monocarboxylate transporters (MCTs). Once in neurons, lactate is oxidized to produce ATP. The ANLS hypothesis is supported by multiple studies, demonstrating a lactate gradient from astrocytes to neurons *in vivo*^15^ or the astrocytic release of lactate upon arousal-induced cortical activity^16^. However, the ANLS theory has faced a long-term controversy^17, 18^, as evidences have arisen that neuronal activation prompts neuronal glucose consumption without lactate uptake^19, 20^, active synapses require glucose consumption to maintain neurotransmitter vesicle recycling and efficient synaptic transmission^12, 21^ and brain rhythms featuring high energy expenditure are only fully sustained by glucose and not lactate^21, 22^.

Despite the importance of this controversy regarding the principles underlying brain energetics, the question of whether brain glucose metabolism can be sustained in the absence of astrocytic glucose transport remains unexplored. Although astrocytes are highly glycolytic and play a key role in the central regulation of energy homeostasis^23–27^ and learning^28–31^, the necessity of GLUT1, i.e. the main astrocytic glucose transporter, for the astrocytic contribution to these processes has never been studied. Here, we aimed to tackle these questions by generating a mouse model lacking GLUT1 specifically in astrocytes and assessing its abilities regarding systemic energy homeostasis and cognition, as well as the molecular and cellular features underlying these competences.

## Results

### GLUT1 is necessary for glucose uptake and metabolism in astrocytes

To elucidate the contribution of GLUT1 to astrocytic glucose uptake and cellular metabolism, we cultured primary astrocytes from *Slc2a1*^flox/flox^ mice and induced GLUT1 deletion via transfection with a Cre-expressing plasmid (GLUT1 KO cells). Transfection efficacy was evaluated by GFP expression observation (Fig. 1a). Successful loss of GLUT1 in astrocytes was confirmed by reductions of both *Slc2a1* mRNA expression (Fig. 1b) and GLUT1 protein levels (Fig. 1c,d) that resulted in a marked reduction of glucose uptake (Fig. 1e). Coherently, GLUT1 KO astrocytes exhibited lower extracellular lactate concentration over 7 days (Fig. 1f) and a diminished lactate release upon stimulation with glucose after starvation (Fig. S1a). Furthermore, real-time extracellular flux analysis revealed that GLUT1 KO astrocytes exhibited reduced glycolytic rate (Fig. 1g, Fig. S1b,c). In view of this glycolytic deficiency, we assessed mitochondrial respiration, finding that GLUT1 KO astrocytes showed intact basal respiration, but failed to reach the same degree of maximal respiration exhibited by control astrocytes (Fig. 1h and Fig. S1d). Noteworthy, GLUT1 KO astrocytes were able to maintain a total ATP production rate comparable to that of control astrocytes (Fig. 1i). Thus, we hypothesized that in basal conditions GLUT1 KO astrocytes are able to meet cellular energy demands using fuels other than glucose. Indeed, GLUT1 KO astrocytes exhibited increased glutamine dependency (Fig. 1j) paired with largely unaltered fatty acid dependency (Fig. S1e). This higher reliance on glutamine oxidation was further corroborated by experiments indicating enhanced glutamine oxidation capacity (Fig. 1k). Overall, these results indicate that GLUT1 is fundamental for astrocytic glucose uptake and metabolism, but not to maintain total ATP production.

**Fig. 1.**
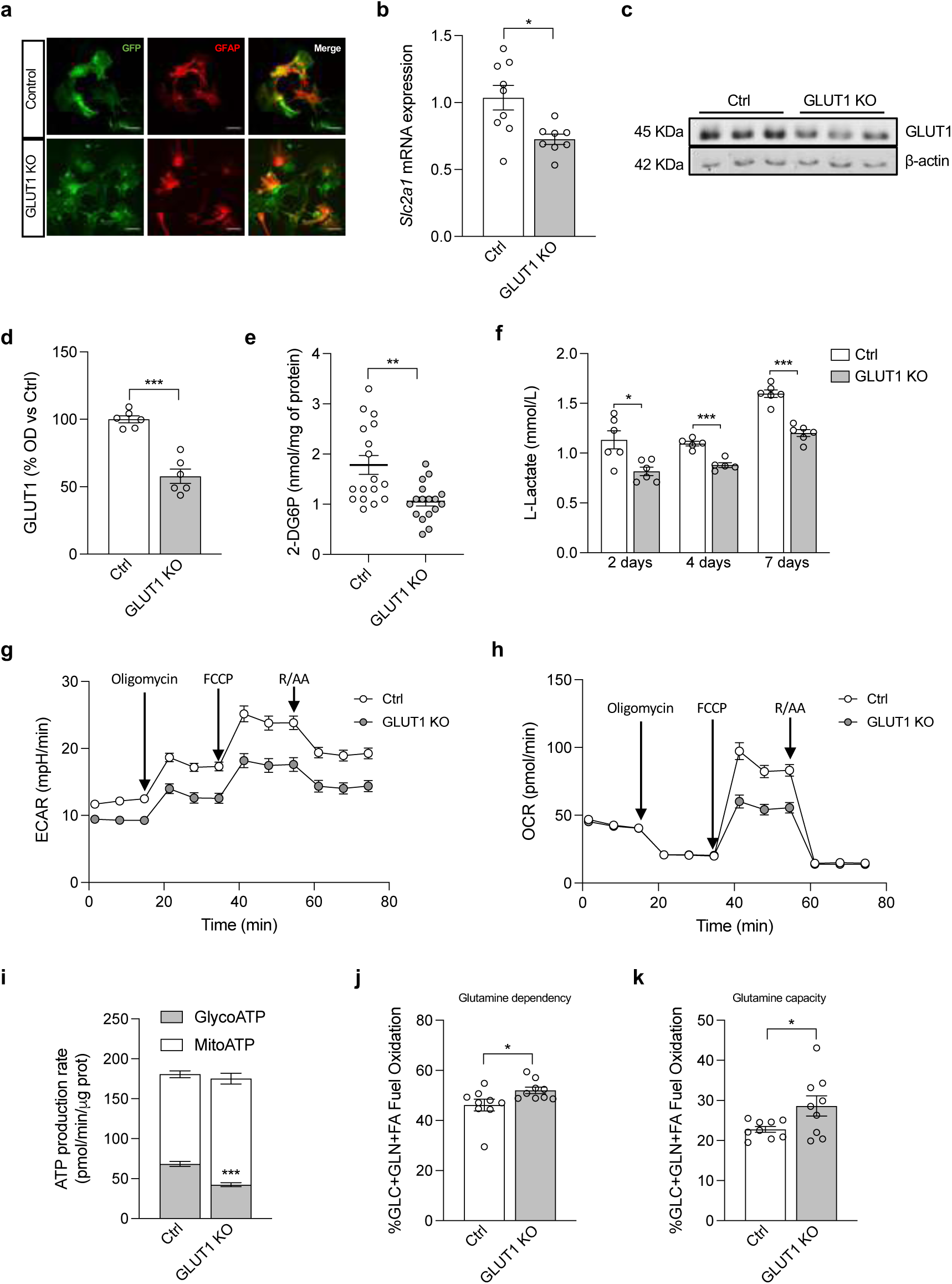
GLUT1 is fundamental for astrocytic glucose uptake and metabolism, but not to maintain total ATP production. **a**, GFP expression in control and GLUT1 KO primary cultured astrocytes (representative image of n = 6 independently isolated astrocyte cultures). Green: GFP; red: GFAP. Scale bar: 50µm. **b,** *Slc2a1* mRNA expression levels in primary cultured astrocytes without (control: Ctrl) or with (GLUT1 KO) Cre recombination (n = 8-9 independently isolated astrocyte cultures). **c,** GLUT1 protein expression level representative western blot image and **d**, quantification (optical density, OD) in primary cultured astrocytes in Ctrl and GLUT1 KO cells (n = 6 independently isolated astrocyte cultures). **e,** Glucose uptake in control and GLUT1 KO cultured astrocytes (n = 17 independent wells). **f,** L-Lactate release in culture medium in control (Ctrl) and GLUT1 KO cultured astrocytes (in mmol/L; n = 6 independently isolated astrocyte cultures). **g,** Glycolytic flux assessed by extracellular acidification rate (ECAR) and **h**, mitochondrial respiration evaluation by oxygen consumption rate (OCR) in Ctrl and GLUT1 KO cultured astrocytes (n = 9 independent wells). **i,** Glycolysis and mitochondrial respiration derived ATP production rate in Ctrl and GLUT1 KO cultured astrocytes (n = 9 independent wells). **j,** Glutamine dependency and **k**, glutamine capacity assessment (n = 9 independent wells). Data are presented as mean ± SEM. *p ≤ 0.05, **p ≤ 0.01, ***p<0.001 as determined by two-tailed Student’s t test in all cases.

### GLUT1 ablation alters astrocytic reactivity and morphology

Given the essential contribution of GLUT1 to astrocytic metabolism shown above, we examined the effects of astrocytic GLUT1 ablation within the mammalian brain (Fig. 2a). To demonstrate that astrocytes underwent Cre recombination and subsequent GLUT1 loss, we purified ACSA-2^+^ cells from the brains of adult mice (Fig. S2a), and confirmed that GLUT1 fluorescent signal (Fig. 2b and Fig. S2b), *Slc2a1* mRNA expression (Fig. 2c) and GLUT1 protein levels (Fig. 2d,e) were significantly decreased in ACSA-2^+^ cells of GLUT1^ΔGFAP^ mice. In this scenario, it is tempting to postulate a potential compensation for the loss of GLUT1 by increased expression of other Slc2a members. Contrarily, the other members of the GLUT gene family showed unaltered mRNA expression upon GLUT1 ablation, as well as a very low mRNA expression compared to that of *Slc2a1* (Fig. S2c).

**Fig. 2.**
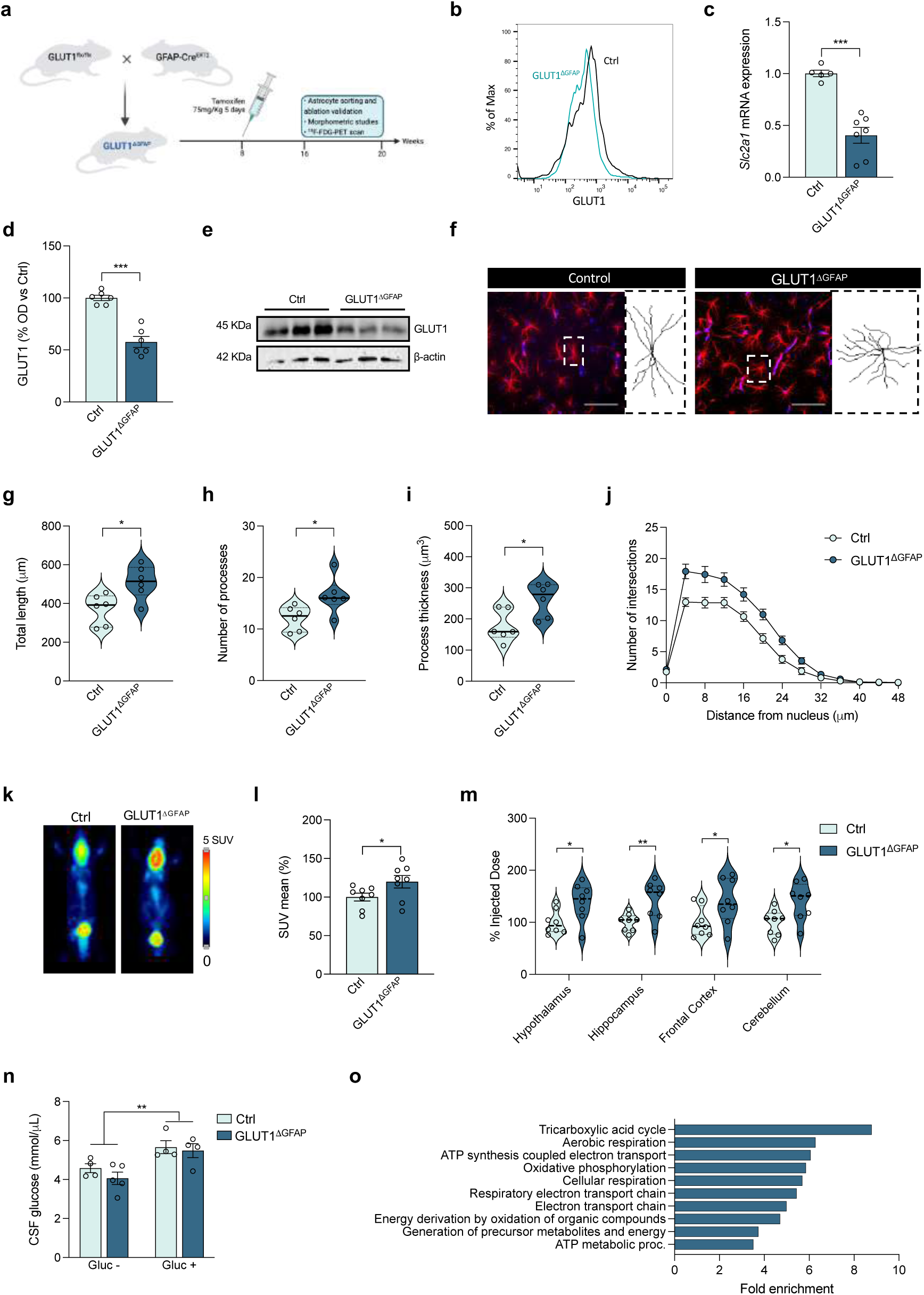
Inducible *in vivo* ablation of astrocytic GLUT1 enhances CNS glucose utilization without altering CSF glucose levels and shifts the whole energetic profile of the brain. **a**, Strategy used to generate mice lacking GLUT1 transporter specifically in astrocytes (GLUT1^1′GFAP^) and their control littermates (GLUT1^f/f^). **b,** GLUT1 fluorescence intensity quantification histogram, **c**, mRNA levels (n = 5-7 mice per group), **d**, protein expression level quantification (optical density, OD) (n = 6 mice per group) and **e**, representative western blotting image in Ctrl and GLUT1^1′GFAP^ mice after ACSA-2^+^ FACS-mediated separation of astrocytes (n = 6 mice per group). **f,** Representative GFAP(red)-DAPI(blue) cell micrograph and its respective reconstruction obtained from SNT of stratum radiatum astrocytes of Ctrl and GLUT1^1′GFAP^ mice (representative image of n = 6 per group). Scale bar 50 µm. **g,** Quantification of processes total length, **h**, number of processes, **i**, GFAP process volume and **j**, Sholl analysis representing the astrocyte complexity (n = 6 animals per group, 10 astrocytes per animal). **k,** PET images of representative mice showing brain ^18^F-FDG signal, **l**, the quantification of positron emission (mean SUV) and **m**, *ex vivo* counting of radioactivity in dissected brain samples (n = 8 mice per group). **n,** CSF glucose presence 30 min after i.p. vehicle or glucose injection in Ctrl and GLUT1^1′GFAP^ mice (n = 4-5 mice per group). **o,** WGCNA “blue module” GO Biological Processes showing significant fold enrichment when comparing control and GLUT1^1′GFAP^ mice (n = 6 per group). Data are presented as mean ± SEM. *p ≤ 0.05, **p ≤ 0.01as determined by two-tailed Student’s t test (c, d, g, h, i, l and m) and two-way ANOVA (n).

Astrocytes undergo a pronounced phenotypic transformation after brain injury and/or disease. In order to evaluate if such a phenotype shift could be also occurring after GLUT1 ablation, we assessed their gene expression profile and morphology. The gene expression analyses showed that GLUT1^ΔGFAP^ astrocytes adopt a distinct molecular state, suggesting higher astrocytic reactivity (Fig. S2d). This genetic signature modification was further accompanied by clear morphological alterations (Fig. 2f). GLUT1^ΔGFAP^ astrocytes display increased total process length (Fig. 2g), increased number of processes (Fig. 2h), and process thickness (Fig. 2i). However, despite the increased process arbor complexity, indicated by the large number of intersections in GLUT1^ΔGFAP^ astrocytes at the same radius (Fig. 2j), the Sholl analysis pointed out that enhanced complexity occurs at the same distance from the soma in GLUT1^ΔGFAP^ astrocytes (Fig. 2j). Taken together, these data indicate that GLUT1 ablation triggers an increase of astrocytic process ramification, but not elongation.

### Astrocyte-specific GLUT1 ablation enhances brain glucose utilization

Despite the privileged position of astrocytes to control glucose access into the brain, whether astrocytic GLUT1 is fundamental for brain glucose metabolism *in vivo* remains unknown. To address this question, we subjected GLUT1^ΔGFAP^ and control mice to ^18^F-fluorodeoxyglucose positron emission tomography (^18^F-FDG-PET) scanning. Strikingly, GLUT1^ΔGFAP^ mice brains exhibited an enhanced ^18^F-FDG-PET signal, not only at the whole-brain level (Fig. 2k and 2l) but also in all individually analysed brain regions (Fig. 2m). In light of this unexpected observation, we assessed whether this phenomenon was paired with increased brain glucose levels. To this end, we examined cerebrospinal fluid (CSF) glucose levels 30 min after i.p. injection of 2g/kg glucose or vehicle (saline), failing to find any differences in CSF glucose level between GLUT1^ΔGFAP^ and control mice (Fig. 2n). These results suggest that astrocytic GLUT1 ablation in adult mice enhances brain glucose metabolism without altering CSF glucose presence.

In an attempt to elucidate the global impact of the enhancement in glucose metabolism observed in GLUT1^ΔGFAP^ mice, we undertook a proteomic profiling of both control and GLUT1^1′GFAP^ brains. We were able to identify 2329 proteins, and we performed weighted gene co-expression network analysis (WGCNA) identifying a total of 27 modules of proteins that were highly co-expressed across our cohorts of mice (Fig. S2e). Only 1 out of the 27 modules (termed “blue module”) exhibited a high correlation with genotype (Fig. S2f-h). Crucially, gene ontology (GO) analysis of the “blue module” revealed that the first ten GO Biological Processes showing a significant fold enrichment were all related to cell energy metabolism (Fig. 2o). Considering that this proteomic profiling was not performed in purified astrocytes but in brain tissue containing all brain cell types, this result suggests that disrupting only astrocytic glucose metabolism is sufficient to shift the whole energetic profile of the brain.

### Astrocytic GLUT1 ablation neither induces brain angioarchitecture alteration nor BBB breakdown

Ablation of GLUT1 in blood-brain-barrier (BBB) endothelial cells induces serious brain angioarchitecture and BBB integrity alterations^32^. As astrocytic endfeet are an essential component of the BBB, we examined whether astrocytic GLUT1 ablation could also induce such alterations. After demonstrating that the Cre/LoxP strategy employed does not induce GLUT1 downregulation in endothelial cells (Fig. S3a-c), our data revealed an absence of serum-borne proteins (IgG and fibrin) in capillary-depleted brains of GLUT1^ΔGFAP^ mice similar to that observed in control mice (Fig. S3d). Besides, we could not find IgG (Fig. S3e) or fibrin (Fig. S3f) leaking out of vascular (lectin^+^) areas, arguing against the possibility of BBB leakage in GLUT1^ΔGFAP^ mice. Consistently, the expression of the gap-junction proteins occludin (Fig. S3g,i,j) and zonulin-1 (ZO-1) (Fig. S3h-j) was also unaltered, supporting the idea of an intact BBB integrity. Furthermore, we examined the brain microvascular structure by quantifying the length of vascular networks. Again, no significant changes were found between GLUT1^ΔGFAP^ and control mice (Fig. S3k) in fractional vascular volume (Fig. S3l) and/or vascular length (Fig. S3m).

### Systemic glucose homeostasis is improved upon astrocyte-specific GLUT1 ablation

To determine whether astrocytic GLUT1 is involved in maintaining whole body glucose homeostasis, we investigated if mice lacking GLUT1 in astrocytes display metabolic alterations. To this end, we performed experiments in mice fed with either a normal chow diet (NCD) or a high-fat diet (HFD), to account for how GLUT1^ΔGFAP^ mice cope with a metabolic challenge. Although no significant differences between genotypes were observed in either body weight (Fig. S4a) or body composition (Fig. S4b), we noted marked differences in feeding behaviour. Specifically, GLUT1^ΔGFAP^ mice showed a greater ability to curb the fasting-elicited hyperphagic response (Fig. 3a), and they exhibited an increased suppression of hyperphagia in response to i.p. glucose administration (Fig. 3b). Furthermore, HFD-fed GLUT1^ΔGFAP^ mice showed an enhanced capacity to readjust systemic glucose levels after i.p. glucose injection-induced hyperglycaemia (Fig. 3c) compared to HFD-fed control mice, without changes in systemic insulin sensitivity (Fig. 3d). Nevertheless, the improved glucose tolerance in HFD-fed GLUT1^ΔGFAP^ mice was coupled with a pronounced increase in glucose-stimulated pancreatic insulin secretion (Fig. 3e).

**Fig. 3.**
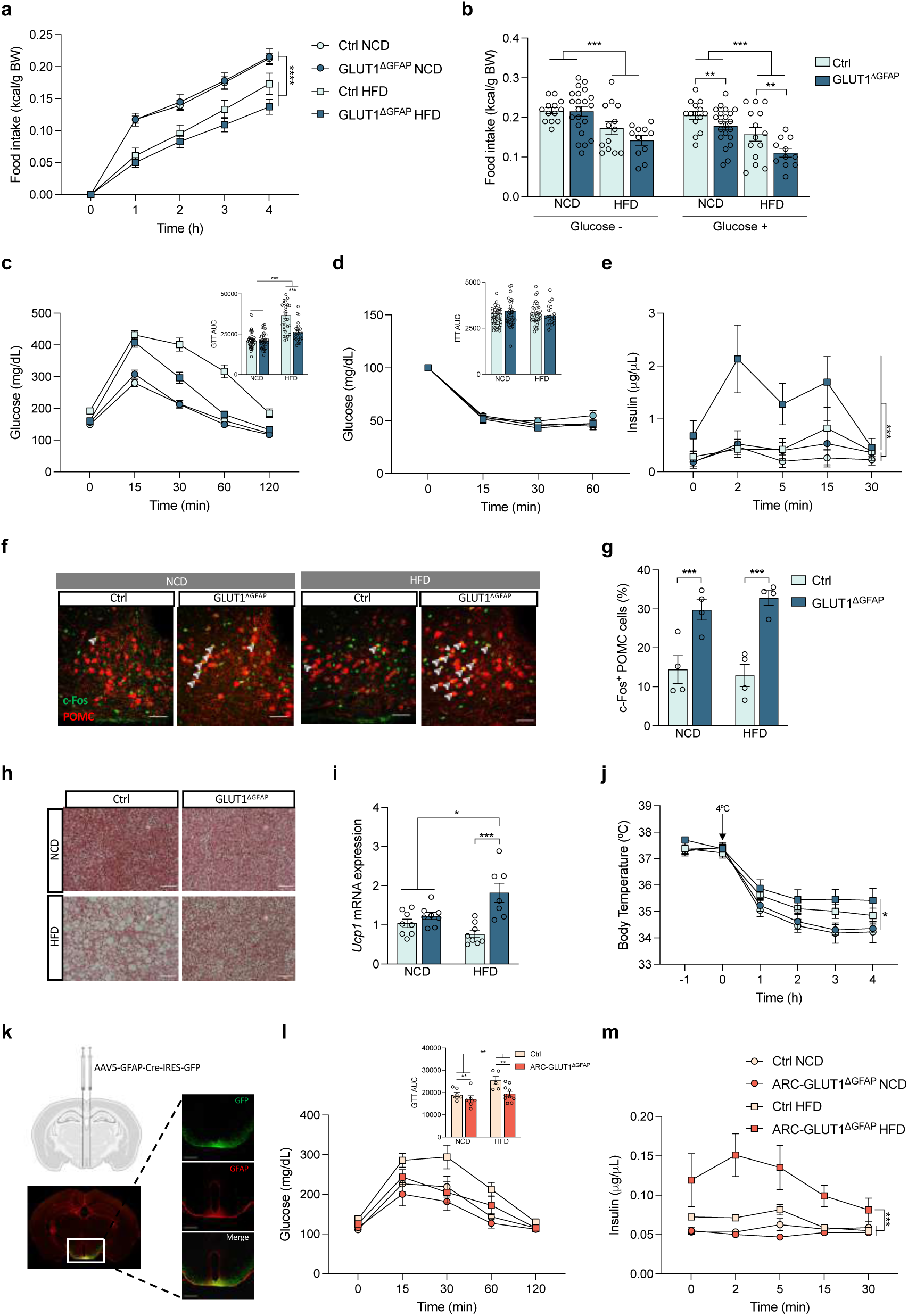
Mice lacking GLUT1 in astrocytes exhibit improved glucose homeostasis. **a**, Fasting-induced hyperphagic response and **b**, 4 h accumulated food intake after i.p. injection with vehicle or glucose in normal chow diet (NCD)- and high fat diet (HFD)-fed control (Ctrl) and GLUT1^1′GFAP^ mice (n = 13-21 per group). **c,** Glucose tolerance test (GTT) and the quantification of the area under the curve (AUC), **d**, insulin tolerance test (ITT) and AUC and **e**, glucose stimulated insulin secretion in NCD- and HFD-fed Ctrl and GLUT1^1′GFAP^ mice (n = 25-40 mice per group in c and d; n = 8 in e). **f,** Representative c-Fos (green) staining of POMC (red) neurons of the arcuate nucleus (ARC) and **g**, quantification of c-Fos^+^-POMC cells in Ctrl and GLUT1^1′GFAP^ mice in NCD or HFD (n = 4 mice per group, 2 slices per mouse). Scale bar 50 µm. **h,** Haematoxylin and eosin staining of brown adipose tissue (BAT) sections in NCD- and HFD-fed Ctrl and GLUT1^1′GFAP^ mice (n = 6 mice per group, 2 slices per mouse). Scale bar 100 µm. **i,** *Ucp1* mRNA expression in Ctrl and GLUT1^1′GFAP^ mice under NCD or HFD (n = 8 mice per group). **j,** Rectal temperature of control and GLUT1^1′GFAP^ mice on NCD and HFD upon cold exposure (+4°C) for 4 h (n = 12-20 mice per group). **k,** Representative illustration and immunostaining image depicting the area infected by AAV5-GFAP-Cre-IRES-GFP (green) and GFAP immunoreactivity (red) (n = 7-10 mice per group). Scale bar 500 µm. **l,** Glucose tolerance test (GTT) and quantification AUC (n = 7-10 mice per group) and **m**, glucose stimulated insulin secretion in NCD- and HFD-fed Ctrl and ARC-specific astrocytic GLUT1 ablated (ARC-GLUT1^1′GFAP^) mice in (n = 4 mice per group). Data are presented as mean ± SEM. *p ≤ 0.05, **p ≤ 0.01, ***p ≤ 0.001 as determined by repeated measurement ANOVA (a, c, d, e, j, l and m) and two-way ANOVA followed by Tukey (b, c AUC, d AUC, g, i and l AUC).

Thus, we analysed the histological features of GLUT1^ΔGFAP^ mice pancreatic islets. We observed that HFD-fed GLUT1^ΔGFAP^ mice islets did not display the hyperplasic features typically shown by HFD-fed mice (Fig. S4c), but were indistinguishable from NCD-fed mice islets (Fig. S4d,e).

The above analysed metabolic features are controlled by glucose-sensing proopiomelanocortin (POMC) neurons in the arcuate nucleus of the hypothalamus (ARC)^33–36^. Noteworthy, it has been shown that astrocytes regulate POMC neuron activation^25, 37^. Hence, we evaluated the activity response of ARC POMC neurons using c-Fos immunoreactivity, uncovering that GLUT1^ΔGFAP^ mice displayed a markedly higher number of c-Fos^+^ POMC cells compared to control animals (Fig. 3f,g).

Brown adipose tissue (BAT) represents a core organ in control of glucose homeostasis. Haematoxylin and eosin (H&E) staining of BAT sections revealed that HFD-fed GLUT1^ΔGFAP^ mice did not present the typical obesity-related increased BAT adiposity as observed in HFD-fed control mice, but displayed a morphology similar to that observed in NCD-fed mice (Fig. 3h). Considering that this phenotype could be correlated with a higher BAT activity, we examined the mRNA expression (Fig. 3i) and protein levels (Fig. S4f-h) of BAT UCP1, finding that both were markedly upregulated in GLUT1^ΔGFAP^ mice, especially when subjected to HFD feeding. Furthermore, we measured BAT activity by determining core body temperature during a cold-exposure, finding that HFD-fed GLUT1^ΔGFAP^ mice were the most efficient group generating heat (Fig. 3j). It is known that glucose-mediated activation of locus coeruleus (LC) tyrosine hydroxylase (TH) neurons is necessary to increase BAT activity upon HFD feeding, preventing obesity^38^. In our hands, the number of c-Fos^+^-TH cells observed in GLUT1^ΔGFAP^ mice was, albeit non-significantly, increased (Fig. S4i,j), potentially contributing to the BAT phenotype and activity observed in those mice. Considering that GFAP is also expressed in other tissues besides the CNS^39^, we next investigated whether ablation of GLUT1 specifically in ARC astrocytes is sufficient to elicit the beneficial effects observed in whole-brain GLUT1^ΔGFAP^ mice. To this end, we used adeno-associated viral (AAV)-mediated Cre recombination to delete GLUT1 specifically in GFAP^+^ astrocytes within the ARC (ARC-GLUT1^ΔGFAP^ mice). First, we confirmed that AAV delivery occurred specifically in the ARC, and that Cre-expressing, AAV-infected cells were GFAP^+^ (Fig. 3k). ARC-specific astrocytic GLUT1 ablation elicited a phenotype that corresponded to that observed in whole-brain GLUT1^ΔGFAP^ mice, exhibiting a tendency towards decreased hyperphagia (Fig. S4k) and improved glucose handling (Fig. 3l), paired with a markedly increased glucose-stimulated pancreatic insulin secretion (Fig. 3m) upon HFD. Interestingly, ARC-GLUT1^ΔGFAP^ did not recapitulate the BAT thermogenesis phenotype (Fig. S4l), indicating that astrocytic GLUT1 deletion in other areas, such as LC, is required for this effect.

### Memory is unaltered in mice lacking GLUT1 in astrocytes

Given that brain glucose metabolism is strongly correlated with cognitive proficiency^40, 41^, we examined the ability of GLUT1^ΔGFAP^ mice to properly establish memory. After excluding locomotor activity alterations (Fig. 4a), we tested recognition memory using the Novel Object Recognition (NOR) task. Both NCD- and HFD-fed GLUT1^ΔGFAP^ mice were as competent as control mice at recognizing the familiar object (Fig. 4b) with a slight improvement of GLUT1^ΔGFAP^ mice in the 24 h task. We then compared the ability of the different mice groups of mice to adequately establish spatial memory in the Morris Water Maze (MWM) task. GLUT1^ΔGFAP^ mice were as efficient as control mice on spatial memory acquisition (Fig. 4c) and memory retention (Fig. 4d). As exposure to a cognitive-demanding training might affect brain glucose metabolism, we subjected all mice groups to ^18^F-FDG-PET scanning immediately after the last MWM probe trial. Our data revealed increased ^18^F-FDG brain signal in MWM-trained control mice compared to naïve control mice, and remarkably, this enhancement was even larger in MWM-trained GLUT1^ΔGFAP^ mice (Fig. 4e,f).

**Fig. 4.**
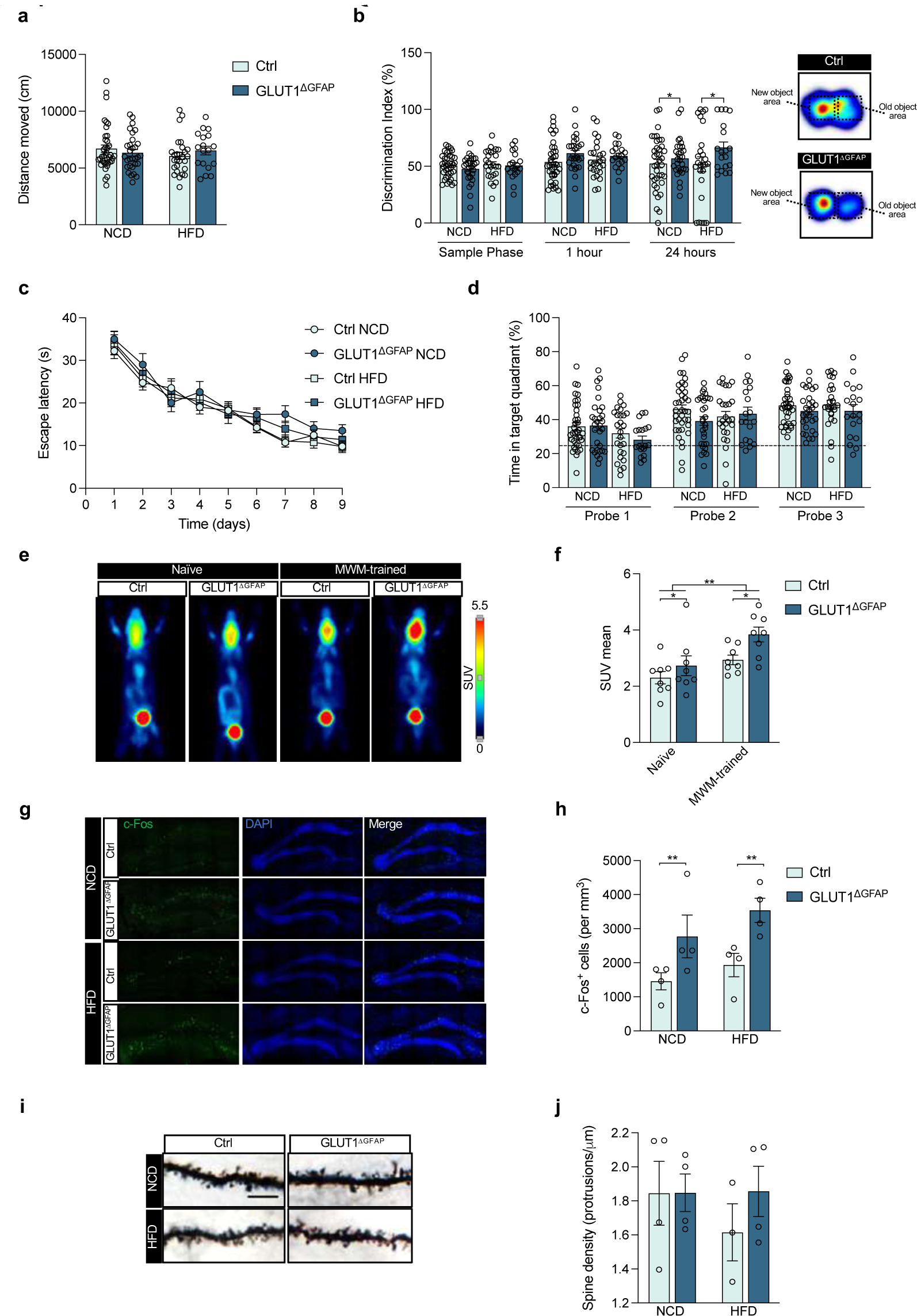
Unaltered memory performance in mice lacking astrocytic GLUT1. **a**, Locomotor activity assessment in NCD- and HFD-fed control (Ctrl) and GLUT1^1′GFAP^ mice (n = 25-40 per group). **b,** Recognition memory assessment by novel object recognition (NOR) task and representative heat map of task performance of Ctrl and GLUT1^1′GFAP^ mice. Data shows discrimination index (n = 20-40 per group). **c,** Spatial memory evaluation by Morris water maze (MWM) acquisition and **d**, retention phase of Ctrl and GLUT1^1′GFAP^ mice in NCD and HFD (n = 20-40 per group). **e,** PET images of representative mice showing brain ^18^F-FDG signal and **f**, the quantification of positron emission (mean SUV) of naïve and MWM-trained Ctrl and GLUT1^1′GFAP^ mice (n = 8 mice per group). **g,** Representative c-Fos (green) staining in the dentate gyrus of hippocampus and **h**, quantification of c-Fos^+^ cells per area in Ctrl and GLUT1^1′GFAP^ mice under NCD or HFD (n = 4 mice per group, 2 slices per mouse). **i,** Representative image of dendritic spines from Golgi-Cox-stained slices and **j**, quantification of the spine density from NCD- and HFD-fed Ctrl and GLUT1^1′GFAP^ mice (n = 4 mice per group, 30 dendrites per animal). Data are presented as mean ± SEM. *p ≤ 0.05, **p ≤ 0.01 as determined by two-way ANOVA followed by Tukey (a, b, d, f, h and j) and repeated measurement ANOVA (c).

Given the slightly improved cognitive abilities upon astrocytic GLUT1 ablation, we aimed to investigate its underlying histological correlates. First, we explored c-Fos immunoreactivity in the dentate gyrus of the hippocampus, finding a markedly higher number of c-Fos^+^ cells in the GLUT1^ΔGFAP^ group compared to controls (Fig. 4g,h). Accordingly, while Golgi-Cox staining revealed no differences between NCD-fed control and GLUT1^ΔGFAP^ mice hippocampal spine density, it evidenced that HFD-fed control mice had a slightly lower spine density, and this effect was reverted in HFD-fed GLUT1^ΔGFAP^ mice (Fig. 4i,j).

### Brain purinergic signalling is essential for improved glucose homeostasis and memory preservation in astrocytic GLUT1-ablated mice

ATP stands among the main bioactive molecules released by astrocytes^42^. According to our data, astrocytes maintain total ATP production rate despite GLUT1 ablation (Fig. 1i). This discovery led us to investigate whether GLUT1-ablated astrocytes also maintained their ability to release ATP. Importantly, GLUT1 KO primary astrocytes featured increased ATP release to the extracellular medium (Fig. 5a) compared to control cells. Consistently, microdialysis-isolated interstitial fluid of freely moving GLUT1^ΔGFAP^ mice presented a larger increase in ATP concentration within the first 30 min after i.p. glucose administration compared to control mice (Fig. 5b). To discard that this effect was not a consequence of the animal stress triggered by manipulation while injecting glucose, a cohort of mice were saline or glucose-injected, demonstrating that saline administration did not induce any raise of ATP levels in the interstitial fluid (Fig. S5a). Therefore, we hypothesized that enhanced brain purinergic signalling could be essential for the systemic glucose homeostasis improvement and memory preservation displayed by GLUT1^ΔGFAP^ mice. Indeed, we found that acute intracerebroventricular (i.c.v.) administration of pyridoxalphosphate-6-azophenyl-2’,4’-disulfonic acid (PPADS) (Fig. 5c), a non-selective P2 purinergic antagonist, totally abrogated both the improved glucose-induced suppression of feeding in NCD (Fig. S5b) or HFD (Fig. 5d) and the enhanced glucose tolerance shown by vehicle-treated GLUT1^ΔGFAP^ mice (NCD Fig. S5c and HFD Fig. 5e). Moreover, PPADS-treated GLUT1^ΔGFAP^ mice exhibited worsened memory in both the NOR (NCD Fig. S5d and HFD Fig. 5f) and MWM tasks particularly on HFD (Fig. 5g-i; data in NCD Fig. S5e,f). Together, these results demonstrate that brain-specific purinergic signalling is necessary for the improved peripheral metabolism and memory persistence phenotypes upon astrocyte-specific GLUT1 ablation.

**Fig. 5.**
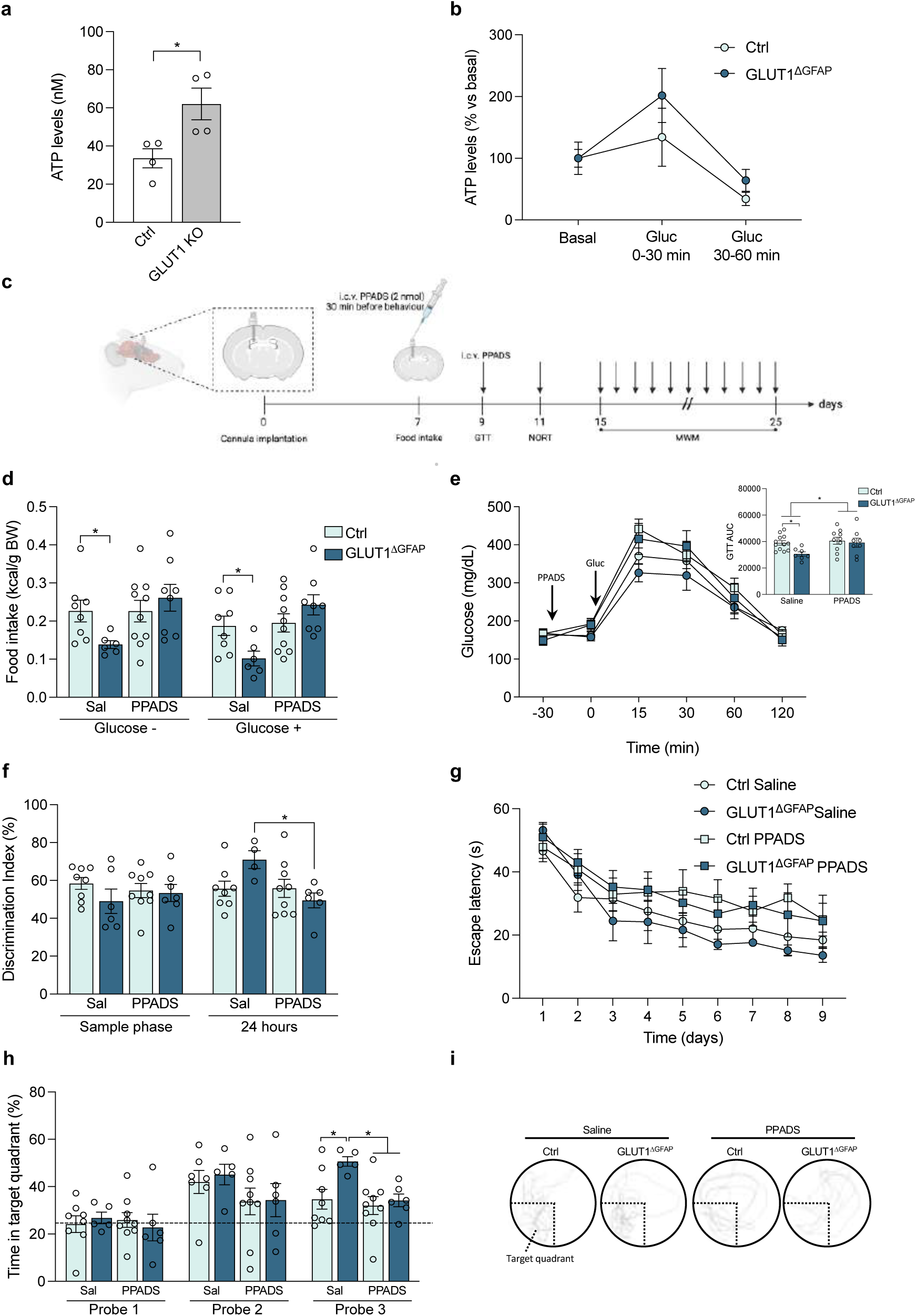
Essential role of brain purinergic signalling in the phenotype exhibited by astrocytic GLUT1-ablated mice. **a**, ATP release from control and GLUT1 KO primary cell cultured astrocytes (n = 4 independently isolated astrocyte cultures). **b,** ATP levels in microdialysis-isolated interstitial fluid of Ctrl and GLUT1^ΔGFAP^ mice before (basal) and after i.p. glucose injection (0-30 and 30-60 min). Data expressed as % of their own basal (n = 8-10 mice per group). **c,** Schematic illustration of intracerebroventricular (i.c.v.) cannulation and PPADS i.c.v. administration 30 min before each metabolic and cognitive assessment task. **d,** Glucose-induced suppression of feeding evaluated by 4 h accumulated food intake in Ctrl and GLUT1^1′GFAP^ mice subjected to HFD after saline or PPADS i.c.v. injection (n = 6-10 per group). **e,** Glucose tolerance test in saline or PPADS i.c.v. treated Ctrl and GLUT1^1′GFAP^ mice on HFD and AUC quantification (n = 8-12 per group). **f,** Recognition memory assessment by novel object recognition (NOR) task of HFD-fed Ctrl and GLUT1^1′GFAP^ mice after saline or PPADS i.c.v. administration (n = 6-9 per group). **g,** Spatial memory evaluation by Morris water maze (MWM) acquisition phase, **h**, retention phase and **i**, a representative image of the swimming path of saline or PPADS i.c.v. treated Ctrl and GLUT1^1′GFAP^ mice on HFD (n = 5-9 per group). Data are presented as mean ± SEM. *p ≤ 0.05 as determined by two-tailed Student’s t test, repeated measurement ANOVA (b, e and g) and two-way ANOVA followed by Tukey (d, e AUC, f and h).

### Astrocytic insulin receptor mediates enhanced ATP release and its ablation impairs glucose homeostasis and memory

Insulin signalling is essential for insulin-dependent ATP release by astrocytes^43^. Intriguingly, we found that FACS-purified astrocytes from GLUT1^ΔGFAP^ mice brains exhibited increased insulin receptor (IR) expression (Fig. 6a). It is thus plausible that astrocyte IR overexpression upon GLUT1 ablation mediates enhanced astrocytic insulin-stimulated ATP release. To test this hypothesis we crossed *Insr*^flox/flox^ (named IR^f/f^) mice with hGFAP-CreER^T2+/-^ in order to obtain the hGFAP-CreER^T2+/-^:IR^f/f^ (named IR^ΔGFAP^) line. Tamoxifen-induced IR ablation was confirmed in sorted astrocytes (Fig. S6a). Crucially, IR^ΔGFAP^ mice failed to increase brain interstitial fluid ATP concentration triggered by i.p. glucose administration (which prompts a physiological insulin release) to the same extent as GLUT1^ΔGFAP^ (Fig. 6b).

**Fig. 6.**
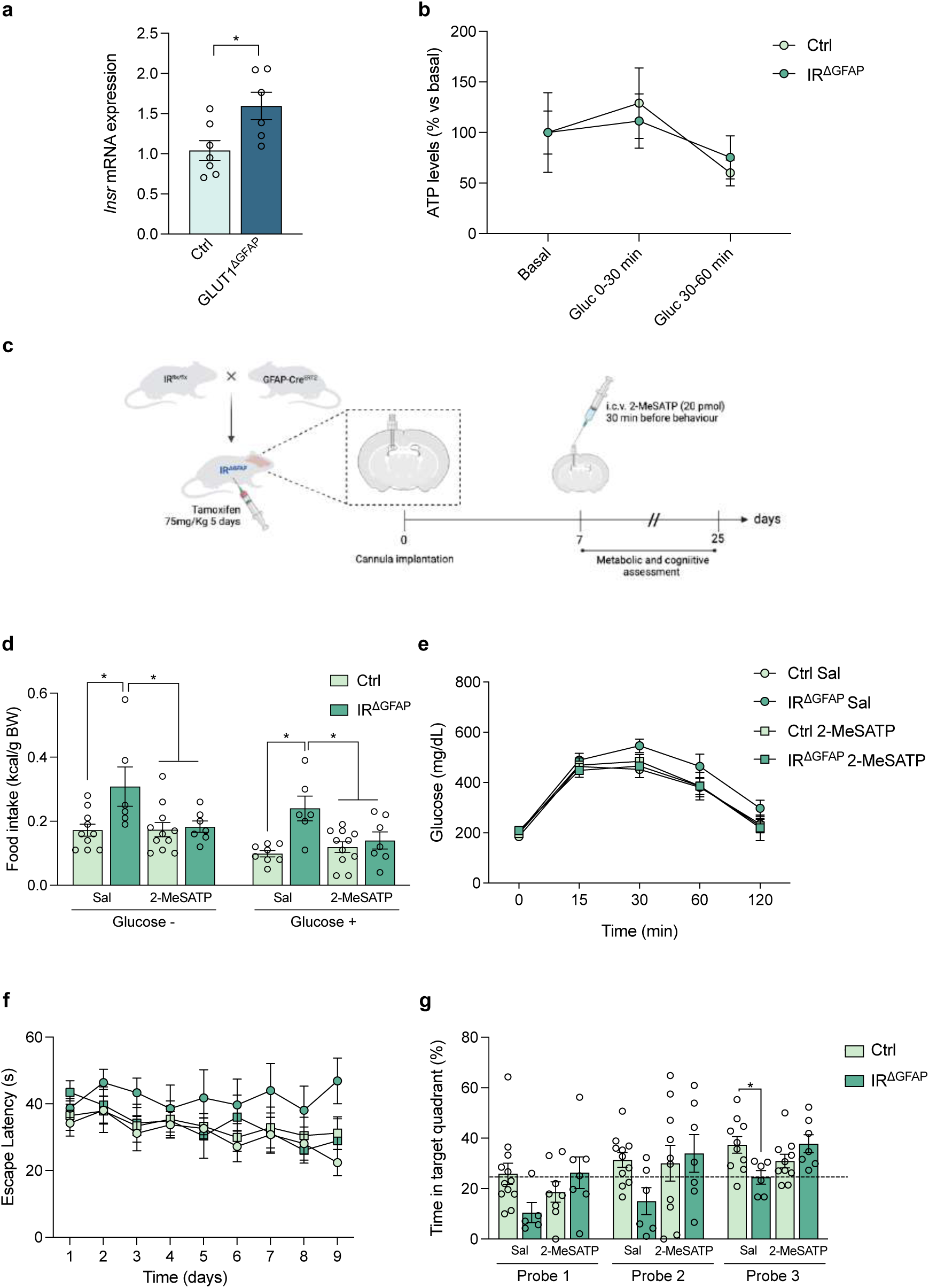
Astrocytic insulin receptor ablation-induced alterations are rescued by purinergic signalling enhancement. **a**, *Insr* (insulin receptor, IR, encoding gene) mRNA levels in control (Ctrl) and GLUT1^1′GFAP^ mice after ACSA-2^+^ FACS-mediated isolation of astrocytes (n = 6-7 mice per group). **b,** ATP levels in microdialysis-isolated interstitial fluid of Ctrl and IR^ΔGFAP^ mice before (basal) and after i.p. glucose injection (0-30 and 30-60 min). Data expressed as % of their own basal (n = 6-8 mice per group). **c,** Schematic illustration of the strategy used to generate mice lacking IR specifically in astrocytes (IR^1′GFAP^) and their control littermates (IR^f/f^), intracerebroventricular (i.c.v.) cannulation and 2-MeSATP i.c.v. administration. **d,** Glucose-induced suppression of feeding evaluated by 4 h accumulated food intake in Ctrl and IR^1′GFAP^ mice subjected to HFD after saline or 2-MeSATP i.c.v. injection (n = 6-11 per group). **e,** Glucose tolerance test in saline or 2-MeSATP i.c.v. treated Ctrl and IR^1′GFAP^ mice on HFD (n = 8-12 per group). **f,** Spatial memory evaluation by Morris water maze (MWM) acquisition and **g**, retention phase of saline or 2-MeSATP i.c.v. treated Ctrl and IR^1′GFAP^ mice on HFD (n = 6-11 per group). Data are presented as mean ± SEM. *p ≤ 0.05 as determined by two-tailed Student’s t test, repeated measurement ANOVA (b, e and f) and two-way ANOVA followed by Tukey (d and g).

To further test the relevance of astrocytic IR for energy homeostasis and cognition, we studied glucose metabolism and memory formation in IR^ΔGFAP^ mice. In agreement with previous studies^25^, we found that IR^ΔGFAP^ mice displayed reduced brain glucose utilization assessed by ^18^F-FDG-PET (Fig. S6b-g) and decreased glucose concentration in the CSF (Fig. S6h). Metabolically, although gaining the same body weight as control mice (Fig. S6i), IR^ΔGFAP^ mice displayed reduced glucose-induced suppression of hyperphagia (Fig. S6j) and lower glucose tolerance (Fig. S6k) in the presence of unaltered insulin sensitivity (Fig. S6l). Provided that brain insulin signalling has an important role in memory and learning^6, 44^, and that its disruption has been linked to cognitive dysfunction^45–47^, we examined recognition and spatial memory in IR^ΔGFAP^ mice. Compared to control mice, although exhibiting identical locomotor activity (Fig. S6m), IR^ΔGFAP^ mice were less efficient memorizing objects in the NOR task (Fig. S6n) and exhibited impaired spatial memory (Fig. S6o,p).

Given the necessity of astrocytic IR for glucose-stimulated increases of brain ATP concentrations demonstrated above (Fig. 6b), we hypothesized that insufficient brain purinergic signalling could be one of the causes underlying the aberrant metabolic and cognitive phenotypes presented by IR^ΔGFAP^ mice. In fact, acute i.c.v. administration of 2-methylthio-ATP (2-MeSATP), a non-selective P2 purinergic agonist (Fig. 6c), was sufficient for IR^ΔGFAP^ mice to recover control mice-like suppression of hyperphagia in NCD (Fig. S6q) and in HFD (Fig. 6d), which was absent in vehicle-treated IR^ΔGFAP^ mice. 2-MeSATP administration also reverted the glucose intolerance shown by IR^ΔGFAP^ mice (NCD Fig. S6r and HFD Fig. 6e). Furthermore, the impairments in spatial memory abilities exhibited by vehicle-treated IR^ΔGFAP^ mice were corrected upon i.c.v. 2-MeSATP treatment in NCD (Fig. S6s,t) and in HFD (Fig. 6f,g). Together, these results provide the first evidence of the relevance of astrocytic IR for memory, and demonstrate that brain-specific purinergic stimulation is sufficient to rescue the aberrant glucose metabolism and cognitive deficits elicited by astrocyte-specific IR ablation.

### Enhanced astrocytic insulin signalling in GLUT1-ablated astrocytes is indispensable for improved glucose homeostasis and memory preservation

In line with above-mentioned evidence of a higher IR expression in GLUT1-deficient astrocytes, an enhanced insulin-stimulated ATP release was observed in cultured GLUT1 KO astrocytes, and this effect was totally abrogated by concomitant treatment with S961, a specific IR antagonist (Fig. 7a). Besides, insulin treatment did not trigger ATP release increase in cultured IR KO astrocytes, confirming the necessity of IR for insulin-stimulated ATP release (Fig. 7a).

**Fig. 7.**
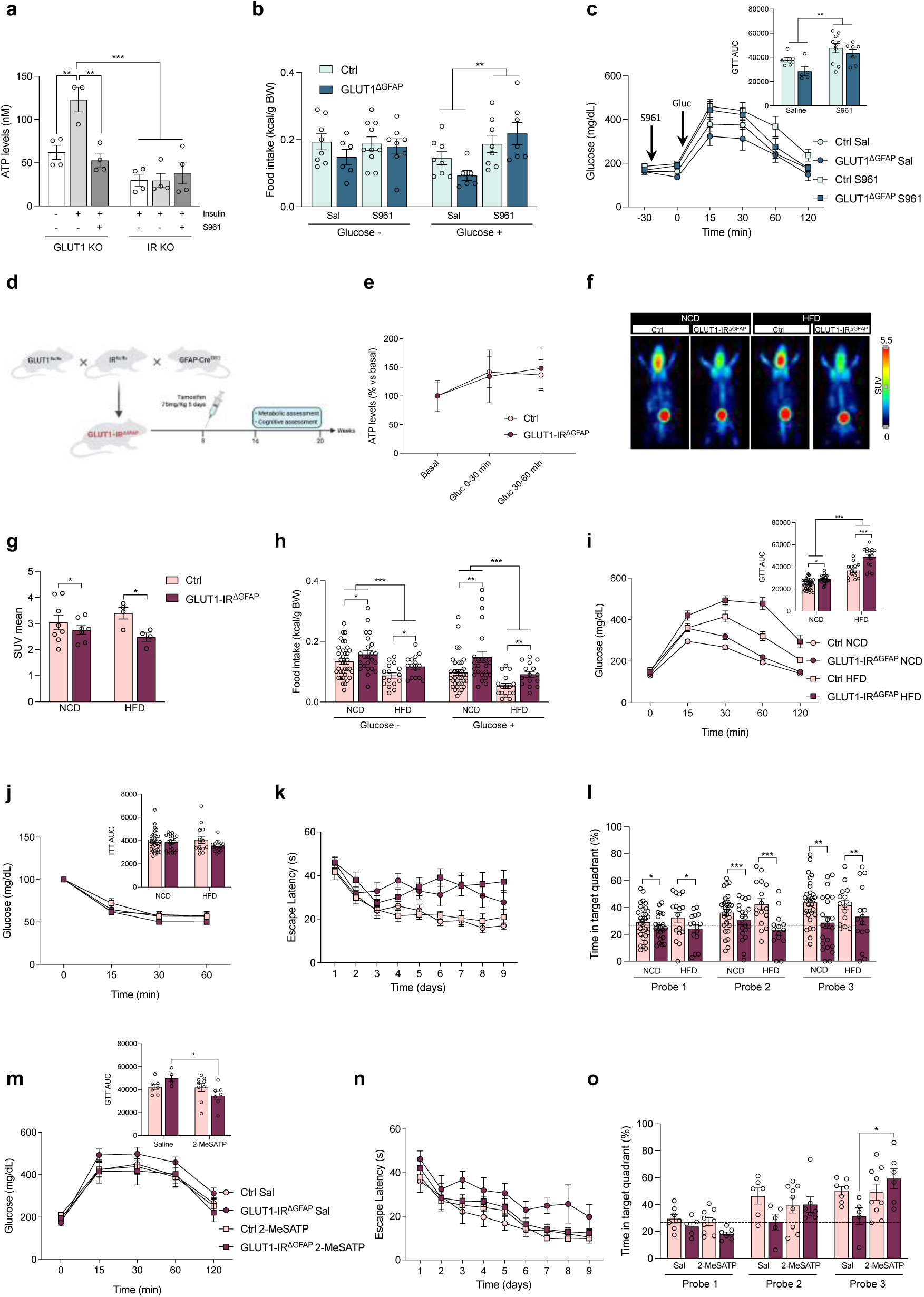
Increased astrocytic insulin receptor signalling is necessary for the ATP signalling-mediated benefits of astrocytic GLUT1 ablation. **a**, ATP release from GLUT1 KO and IR KO astrocytes exposed to either vehicle, insulin or insulin + S961 (insulin receptor antagonist) (n = 4 independently isolated astrocyte cultures). b, Glucose-induced suppression of feeding evaluated by 4 h accumulated food intake in control (Ctrl) and GLUT1^1′GFAP^ mice subjected to HFD after saline or S961 i.c.v. injection (n = 6-10 per group). c, Glucose tolerance test in saline or S961 i.c.v. treated Ctrl and GLUT1^1′GFAP^ mice on HFD and the corresponding AUC quantification (n = 6-10 per group). d, Strategy used to generate mice concomitantly lacking GLUT1 and IR specifically in astrocytes (GLUT1-IR^1′GFAP^) and their control littermates (GLUT1-IR^f/f^). e, ATP levels in microdialysis-isolated interstitial fluid of Ctrl and GLUT1-IR^ΔGFAP^ mice before (basal) and after i.p. glucose injection (0-30 and 30-60 min) (n = 8 mice per group). f, PET images of representative mice showing brain ^18^F-FDG signal and g, the quantification of positron emission (mean SUV) in Ctrl and GLUT1-IR^1′GFAP^ mice under NCD or HFD (n = 4-8 mice per group). h, Glucose-induced suppression of feeding, i, glucose tolerance test with its corresponding AUC and j, insulin tolerance test with its AUC in NCD- and HFD-fed control and GLUT1-IR^1′GFAP^ mice (n = 15-30 per group). k, Spatial memory evaluation by Morris water maze (MWM) acquisition and l, retention phase of Ctrl and GLUT1-IR^1′GFAP^ mice on NCD and HFD (n = 15-30 per group). m, Glucose tolerance test in saline or 2-MeSATP i.c.v. treated control and GLUT1-IR^1′GFAP^ mice on HFD (n = 5-9 per group). n, Spatial memory evaluation by MWM acquisition and o, retention phase of saline or 2-MeSATP i.c.v. treated control and GLUT1-IR^1′GFAP^ mice on HFD (n = 5-9 per group). Data are presented as mean ± SEM. *p ≤ 0.05, **p ≤ 0.01, ***p ≤ 0.001 as determined by two-way ANOVA followed by Tukey (a, b, g, h, i AUC, j AUC, l, m AUC and o) and repeated measurement ANOVA (c, e, i, j, k, m and n).

Taking all these data into consideration, we examined whether insulin signalling is necessary for the metabolic and cognitive improvements observed in GLUT1^ΔGFAP^ mice. Acute i.c.v. S961 administration blunted the improved fasting-induced hyperphagia (NCD Fig. S7a and HFD Fig. 7b) and enhanced glucose tolerance found in GLUT1^ΔGFAP^ mice (NCD Fig. S7b and HFD Fig. 7c). These results evidence that GLUT1^ΔGFAP^ mice require fully operative brain insulin signalling to display their improved systemic glucose homeostasis. However, considering that S961 can antagonize the IR present in most brain cell types^10^, these data are unable to differentiate the specific contribution of astrocytic IR to the phenotype of GLUT1^ΔGFAP^ mice. To address this relevant question, we crossed IR^ΔGFAP^ with GLUT1^ΔGFAP^ mice to generate hGFAP-CreER^T2+/-^:*Slc2a1*^f/f^:*Insr*^f/f^ (named GLUT1-IR^ΔGFAP^) mice, in which tamoxifen administration induces concomitant ablation of GLUT1 and IR exclusively from astrocytes (Fig. 7d). Firstly, microdialysis experiments showed that GLUT1-IR^ΔGFAP^ failed to increase brain ATP levels upon i.p. glucose injection (Fig. 7e) to the same extent as GLUT1^ΔGFAP^ mice. Moreover, ^18^F-FDG-PET studies revealed that GLUT1-IR^ΔGFAP^ mice not only lacked the characteristic enhanced ^18^F-FDG signal present in GLUT1^ΔGFAP^ mice, but featured diminished brain glucose utilization similar to what is observed upon isolated IR deficiency (Fig. 7f,g and Fig. S7c-f). Besides, concomitant ablation of IR and GLUT1 in astrocytes, without inducing body weight differences (Fig. S7g), abolished the beneficial effects regarding both food intake (Fig. 7h) and glucose tolerance observed in GLUT1^ΔGFAP^ mice (Fig. 7i), without altering insulin sensitivity (Fig. 7j). Finally, GLUT1-IR^ΔGFAP^ mice failed to maintain the spatial memory acquisition (Fig. 7k) and retention (Fig. 7l) observed in GLUT1^ΔGFAP^ mice, in absence of locomotor activity alterations (Fig. S7h). Provided that the effects of astrocytic IR ablation were rescued by acute i.c.v. administration of the 2-MeSATP, we decided to treat GLUT1-IR^ΔGFAP^ mice with this same compound. Crucially, i.c.v. administration of 2-MeSATP was sufficient to revert both the inability to properly control peripherally-induced hyperglycaemia (NCD Fig. S7i and HFD Fig. 7m) and the memory deficits shown by GLUT1-IR^ΔGFAP^ mice (NCD Fig. S7j,k; HFD Fig. 7n,o).

Taken together, these results indicate that enhanced IR expression in GLUT1^ΔGFAP^ mice is a necessary feature to improve glucose homeostasis and to maintain memory. In addition, these results also show that brain-specific purinergic stimulation is sufficient to rescue the metabolic and cognitive deficits induced upon concomitant GLUT1 and IR ablation in astrocytes.

## Discussion

It is well established that GLUT1 is the transporter mediating glucose access across the BBB into the brain^13, 48, 49^. Particularly, the lethality of mice with inducible ablation of GLUT1 in the BBB vascular endothelial cells has highlighted the crucial role of endothelial GLUT1 expression in control of brain glucose uptake and energetics^50, 51^. Astrocytic GLUT1-containing endfeet cover almost the entire surface of the blood vessels^52^ with over than 99% of the astrocytes being connected to blood vessels^53^. Given these features and the well-accepted ANLS hypothesis, according to which astrocytes must metabolize glucose and export lactate to neurons for the latter to meet energy needs^14^, one could hypothesize that ablating GLUT1, the main glucose transporter expressed in these cells, would lead to a breakdown of brain glucose uptake and metabolism. Against these expectations, our data indicate that astrocytic GLUT1 ablation results in a paradoxical enhancement of brain glucose utilization. In light of the decreased glycolysis observed upon GLUT1 deletion in cultured astrocytes together with the finding that astrocytic GLUT1 ablation *in vivo* is not compensated by increased expression of any other member of the GLUT family, it is tempting to speculate that other brain cell types could be responsible for the increased glucose metabolism observed in GLUT1^ΔGFAP^ mice. This idea has been already suggested by modelling studies predicting that the majority of glucose diffuses from endothelial cells independently of astrocytic endfeet, and throughout the extracellular fluid to more distant brain cells, facilitating rapid GLUT3-mediated uptake into neurons^13^. The validity of this model has been elegantly proven *in vivo* showing that upon stimulation, glucose consumption by neurons is dramatically increased shifting to a highly glycolytic profile to account for the elevated energy demand^20^. Indeed, our discoveries that brain glucose metabolism is not simply maintained, but enhanced in GLUT1^ΔGFAP^ mice, and that a network of energy metabolism-related proteins is shifted in our proteomic profiling indicates that astrocytic GLUT1 is important enough to trigger a reprogramming of the overall brain metabolism sufficient to sustain energy requirements upon its ablation.

Moreover, mice lacking GLUT1 in astrocytes exhibited improved abilities regarding glucose homeostasis. Glucose-sensing POMC neurons in the ARC are well established as central mediators of satiety^54^, being also able to regulate insulin secretion^36^ and glucose metabolism^34, 55^. Remarkably, POMC neuron glucose sensitivity becomes impaired in HFD-induced obesity, losing their glucose homeostasis regulatory properties^55^. Likewise, glucose-sensing TH neurons in the LC increase body energy expenditure during HFD-induced obesity by prompting BAT activation^38^. Very interestingly, our data reveal that particularly, HFD-fed animals benefit from astrocytic GLUT1 ablation, exhibiting enhanced glucose homeostasis in comparison to HFD-fed control animals. Provided that the loss of GLUT1 specifically in astrocytes of HFD-fed animals induced enhanced glucose-stimulated activation of both ARC POMC neurons and LC TH neurons, this leads us to argue that astrocytic GLUT1 ablation ameliorated the glucose sensing of this neuronal populations, highlighting the importance of these neuronal populations in the activation of peripheral processes that facilitate glucose metabolism such as insulin secretion or BAT activity, respectively.

Brain glucose metabolism and cognitive proficiency are intimately correlated^40, 41^. Although it has been established that astrocytic lactate is needed for memory formation^31^, and that BBB endothelial cell GLUT1 ablation triggers cognitive breakdown^32^, the necessity of astrocytic GLUT1 for cognitive performance has never been tested. Strikingly, far from experiencing a cognitive breakdown, mice lacking GLUT1 in astrocytes were able to form and retain memories, even showing both more active hippocampal neurons and cognitive task-elicited brain glucose metabolism in PET studies compared to control mice. All of these features can be considered as a readout of healthy memory neurocircuits, arguing against the presupposition that astrocytic GLUT1 is as necessary as endothelial GLUT1 to fuel cognitive abilities, and against the necessity of astrocytic glycolysis for memory formation.

In light of the metabolic and cognitive improvement observed, we found that GLUT1 ablation resulted in an enhanced capacity of astrocytes to release ATP. It has been described that IR activation by insulin in astrocytes mediates ATP exocytosis^43^, and we found elevated IR levels in GLUT1-ablated astrocytes, which were responsible for maintaining astrocytic insulin-dependent ATP release. Thus, both the combination of enhanced glucose-stimulated insulin release and astrocytic IR overexpression may cooperate to elicit higher ATP release. This model is further supported by our data indicating that the beneficial effect of astrocyte GLUT1 deletion is reversed by both inhibition of purinergic signalling and simultaneous IR ablation, the effect of which can be overcomed by activation of purinergic signalling.

Collectively, this study uncovered a central role of astrocytes orchestrating brain energy metabolism. Specifically, astrocytic GLUT1 ablation induces an “astrocytic glucose scarcity” situation with lower astrocytic glucose uptake and glycolysis. In this scenario, a fundamental astrocytic reprogramming is conducted with IR overexpression and improved ATP release. Further, the enhanced purinergic signalling could propagate direct or indirect signals to neurons indicating the necessity of a higher metabolic rate (that could be obtained from higher glucose metabolism) and inducing its activity which is translated into improved peripheral energy metabolism and preserved cognition.

## Acknowledgements

We acknowledge all JPOST Team for helping with the mass spectrometric data deposit in ProteomeXChange/PRIDE. We thank excellent contributions from the micro-PET core facility of the University of Navarra (Maria Collantes and Marga Ecay) and the Morphology Core Facility (Universidad de Navarra, Centro de Investigación Médica Aplicada -CIMA-) for the support with histological studies. We thank Dr. Amanda Sierra for her generous protocol sharing and support. This work has been supported by the Spanish Ministry of Economy and Competitiveness (SAF2017-87619-P and PID2021-128737NB-I00, Spanish Government). This work was also supported by a NARSAD Young Investigator grant from the Brain & Behavior Research Foundation (Grant number 28177). C.G.A. is supported by an FPU (Formación de Profesorado Universitario) fellowship from the Spanish Ministry of Universities (FPU18/01458).

## Author contribution

M.S. and C.G.A. conceived the project, designed the experiments, conducted the experiments, analysed the data, and wrote the manuscript with input from the other authors. A.d.C. contributed in behavioural experiments. J.F-I. and E.S designed and performed proteomics studies. M.E-H. analysed proteomic data. E.P. and J.E.O. contributed in microdialysis experiments. A.U., C.E., M.A., C.O-S. designed, performed and analysed CUBIC experiments. G.K. and J.C.B. provided GLUT1^flox/flox^ animals. M.S. and C.G.A. wrote the manuscript. M.J.R. and J.C.B. reviewed the first draft. All authors discussed the data, commented on the manuscript before submission, and agreed with the final submitted manuscript.

## Methods

### Genetic Mouse Models

#### GLUT1^f/f^ Mice

*Slc2a1*^flox/flox^ (named GLUT1^f/f^) mice^56^ were kindly provided by Jens C. Brüning (Max Planck Institute for Metabolism Research, Cologne, Germany), with the permission of Gérard Karsenty (Columbia University, New York, USA).

#### IR^f/f^ Mice

*Insr*^flox/flox^ (named IR^f/f^) mice^57^ were purchased from The Jackson Laboratory (B6.129S4(FVB)-Insr^tm1Khn^/J; Strain 006955; RRID:IMSR_JAX:006955).

#### hGFAP-CreER^T^^2^ Mice

hGFAP-CreER
^T^^2^ mice^58^ were purchased from The Jackson Laboratory (B6.Cg-Tg(GFAP-cre/ERT2)505Fmv/J; Strain 012849; RRID:IMSR_JAX:012849).

#### GLUT1^1′GFAP^ Mice

hGFAP-CreER^T2^ mice were mated with GLUT1^f/f^ mice to obtain hGFAP-CreER^T2^:GLUT1^f/f^ mice (named GLUT1^1′GFAP^), and subsequently breeding colonies were maintained by mating GLUT1^f/f^ mice with hGFAP-CreER^T2^:GLUT1^f/f^ mice. The hGFAP-CreER^T2^ allele was maintained in heterozygous conditions whereas the floxed GLUT1 allele was maintained in homozygous conditions. Animals were kept on a C57BL/6J background. Cre-dependent ablation of the floxed GLUT1 allele was accomplished by tamoxifen (Sigma-Aldrich, T5648) administration (75mg/kg i.p. every 24h for 5 consecutive days) at 8 weeks of postnatal age, as described below (see “Tamoxifen Treatment”).

#### IR^1′GFAP^ Mice

hGFAP-CreER^T2^ mice were mated with IR^f/f^ mice to obtain hGFAP-CreER^T2^:IR^f/f^ mice (named IR^1′GFAP^), and subsequently breeding colonies were maintained by mating IR^f/f^ mice with hGFAP-CreER^T2^:IR^f/f^ mice. The hGFAP-CreER^T2^ allele was maintained in heterozygous conditions whereas the floxed IR allele was maintained in homozygous conditions. Animals were kept on a C57BL/6J background. Cre-dependent ablation of the floxed IR allele was accomplished by tamoxifen (Sigma-Aldrich, T5648) administration (75mg/kg i.p. every 24h for 5 consecutive days) at 8 weeks of postnatal age, as described below (see “Tamoxifen Treatment”).

#### GLUT1-IR^1′GFAP^ Mice

hGFAP-CreER^T2^:GLUT1^f/f^ mice were mated with IR^f/f^ mice to obtain hGFAP-CreER^T2^:GLUT1^f/f^:IR^f/f^ mice (named GLUT1-IR^1′GFAP^), and subsequently breeding colonies were maintained by mating GLUT1^f/f^:IR^f/f^ mice with hGFAP-CreER^T2^:GLUT1^f/f^:IR^f/f^ mice. The hGFAP-CreER^T2^ allele was maintained in heterozygous conditions whereas the floxed GLUT1 and floxed IR alleles were maintained in homozygous conditions. Animals were kept on a C57BL/6J background. Cre-dependent ablation of the floxed GLUT1 and IR alleles was accomplished by tamoxifen (Sigma-Aldrich, T5648) administration (75mg/kg i.p. every 24h for 5 consecutive days) at 8 weeks of postnatal age, as described below (see “Tamoxifen Treatment”).

### Genotyping of Mouse Models

Genotyping of loxP-flanked alleles was performed by PCR using the following custom-designed primer sets, designed to bind to *Slc2a1*^flox/flox^: (Fwd) 5’-TTG AGA GCC ATC TGG AAG GGG G -3’ and (Rev) 5’-CAA GAC TCT GAG GAT GGT GGC CA-3’; or designed to bind to *Insr*^flox/flox^: (Fwd) 5’-GGG GCA GTG AGT ATT TTG GA-3’ and (Rev) 5’-TGG CCG TGA AAG TTA AGA GG-3’. Genotyping of hGFAP-CreER^T2^ was performed by PCR using the following custom-designed primer sets, designed to bind to CreER^T2^: (Fwd) 5’-GCC AGT CTA GCC CAC TCC TT-3’ and (Rev) 5’-TCC CTG AAC ATG TCC ATC AG-3’. For hGFAP-CreER^T2^ genotyping the following internal controls were employed: (oIMR7338) 5’-CTA GGC CAC AGA ATT GAA AGA TCT-3’ and (oIMR7339) 5’-GTA GGT GGA AAT TCT AGC ATC ATC C-3’. All genotyping experiments were performed using the KAPA Hotstart Mouse Genotyping Kit (KAPA Biosystems, KK7352) according to manufacturer’s instructions.

### Tamoxifen Treatment

To excise loxP sites by Cre recombination, 8-week-old hGFAP-CreER^T2^:GLUT1^f/f^, hGFAP-CreER^T2^:IR^f/f^, and hGFAP-CreER^T2^:GLUT1^f/f^:IR^f/f^ mice were subjected to tamoxifen administration. Briefly, tamoxifen (Sigma-Aldrich, T5648) was dissolved at 55°C in a 10% ethanol – 90% peanut oil (Sigma-Aldrich, P2144) solution at a concentration of 20mg/mL. This solution was used to inject mice 75mg/kg of tamoxifen i.p. daily for 5 consecutive days. In order to maintain the same experimental conditions, all the corresponding control groups (GLUT1^f/f^, IR^f/f^ and GLUT1^f/f^:IR^f/f^) also received the tamoxifen treatment.

### Animal Care

All animal procedures were conducted in compliance with the European and Spanish regulations (2003/65/EC; 1201/2005) for the care and use of laboratory animals, and approved by the Ethical Committee of University of Navarra (ethical protocol number 076-19). Animals were housed in groups of 3-7 at 22-24°C, using a 12-h light/12-h dark cycle. Mice had *ad libitum* access to water and the prescribed diet at all times, and food was only withdrawn if required for an experiment. After weaning, mice were fed either on a normal chow diet (NCD) containing 67% calories from carbohydrates, 20% from protein and 13% from fat (Teklad Global 14% Protein Diet, Envigo, 2014) or on a high-fat diet (HFD) containing 20% calories from carbohydrates, 20% from protein and 60% from fat (Research Diets, D12492) as indicated. Experiments were conducted in adult mice (16-24 weeks), using both male and female mice. Mice were randomly assigned to diet and/or treatment experimental groups.

### Primary Mouse Astrocyte Cultures

#### Astrocyte Isolation and Culture

Astrocytes from the hippocampi, hypothalami and cortices were isolated from GLUT1^f/f^ or IR^f/f^ mice pups at postnatal day 0-4 as follows: hippocampi, hypothalami and cortices were isolated in DMEM-GlutaMAX (GIBCO, Thermo Fisher Scientific, 61965-026) and meningeal fragments were carefully removed. Isolated brain areas were transferred to a 15 mL Falcon tube with the dissection media, and were mechanically disaggregated by applying gentle pressure and suction with a disposable pipette. Thereafter, the tube was filled with pre-warmed HBSS and subjected to centrifugation (1 min, 6000 rpm). Supernatant was discarded, and the remaining pellet was resuspended in 10 mL of disaggregation solution (8.5 mL HBSS (GIBCO, Thermo Fisher Scientific, 14170-088), 0.5 mL HEPES 1M (Sigma-Aldrich, H0887), 0.25 mL trypsin 10 mg/mL (Sigma-Aldrich, T9935) and 0.5 mL DNase I 10 mg/mL (Roche, 10104159001) and incubated 15 min at 37°C. Disaggregation was then terminated by addition of 0.25 mL trypsin inhibitor (GIBCO, Thermo Fisher Scientific, 17075-029) at 10 mg/mL and 100 µL of 1% BSA followed by 5 mL of complete culture medium (DMEM-GlutaMAX supplemented with 10% heat-inactivated fetal bovine serum (FBS; GIBCO, Thermo Fisher Scientific, 10500-064) and 0.2% both penicillin 100 IU/ml and streptomycin 100 μg/ml (Lonza Bioscience, DE17-602E). Subsequently, samples were centrifuged (10 min, 6000 rpm) and the supernatant was discarded. The resulting pellet was resuspended in complete culture medium, and placed in a T75 flask (pre-coated with 50 μg/mL poly-D-lysine (Merck Millipore, A-003-E) for 1 hour). Cells were allowed to grow subjected to 5% CO_2_ at 37°C for one week. After this time, primary astrocytes were purified using the McCarthy and de Vellis method^59^. Briefly, primary glial culture-containing T75 flasks were subjected to a 12-hour shaking at 250 rpm, maintained at 5% CO_2_ and 37°C. This shaking was sufficient to detach all brain cell types (microglia and oligodendrocyte precursor cells), but not astrocytes, which remained attached to the T75 internal surface. Detached cell-containing medium was removed, and the remaining cells were subjected to two washings with phosphate buffer saline (PBS; GIBCO, Thermo Fisher Scientific, 14190-094), followed by the addition of fresh complete culture medium. Astrocyte purity of the cultures was assessed by checking the number of GFAP-positive cells by immunofluorescence in a subset of cells, and only cultures in which GFAP-positive cells were higher than 95% were used to perform experiments.

#### Astrocyte Transfection and Selection

Astrocytes were detached from the T75 flask using TrypLE^TM^ (GIBCO, Thermo Fisher Scientific, 12605-028) and were seeded in two 6-well culture plates, subsequently being subjected to plasmid transfection when cell confluence reached 75%. Briefly, 4μg of either control plasmid (“pCMV-Myc-GFP” plasmid, Addgene, 83375) or pCMV-Cre-GFP plasmid (“Cre Shine” plasmid, Addgene, 37404) were added to 2.5% Lipofectamine^TM^-2000 (Thermo Fisher Scientific, 11668027) in Opti-MEM^TM^ (GIBCO, Thermo Fisher Scientific, 31985047) and incubated at room temperature (RT) for 15-20 min. Cells were washed with PBS and 1 mL of the plasmid-Opti-MEM-Lipofectamine solution was added to each well. Plates were incubated for 2 hours subjected to 5% CO_2_ at 37°C. Thereafter, the plasmid-containing solution was removed and cells were extensively washed with HBSS, and cultured in complete culture medium. Both pCMV-GFP and pCMV-Cre-GFP plasmids incorporate a neomycin-resistance (NeoR) sequence. Therefore, 2 days after transfection, positive selection of transfected astrocytes was performed by treating cells for 36 h with Geneticin^®^ (G-418 Sulfate; Thermo Fisher, 11811023) 100 μg/mL in complete culture medium. After extensive washing in HBSS, cells were incubated in complete culture medium for 3 additional days before being subjected to final experiments.

pCMV-Myc-GFP^60^ was a gift from Wesley Sundquist (Addgene plasmid, 83375; http://n2t.net/addgene:83375; RRID:Addgene_83375). Cre Shine was a gift from Thomas Hughes (Addgene plasmid, 37404; http://n2t.net/addgene:37404; RRID:Addgene_37404).

### Real-Time Cell Metabolic Analysis

Extracellular metabolic flux dynamics were measured in primary astrocytes using the XFp Extracellular Flux Analyser (“Seahorse”, Agilent Technologies). The day prior to the assay, astrocytes were seeded in XFp plates at a density of 20000 cells/well and maintained in complete culture medium for 24 h. Thereafter, 1 h prior to the assay, cells were transferred to Seahorse XF DMEM medium (Agilent Technologies) supplemented with 10 mM glucose, 2 mM glutamine and 1 mM pyruvate, and were kept in this medium during the whole assay. All results were normalized by protein content, which was quantified using the Bio-Rad Protein Assay (Bio-Rad) according to manufacturer’s instructions.

#### Mitochondrial Respiration and Glycolysis

Mitochondrial respiration-derived oxygen consumption rate (OCR) and glycolysis-derived extracellular acidification rate (ECAR) were measured in primary astrocytes under basal conditions and in response to 1 μg/mL oligomycin, 1 μM fluoro-carbonyl cyanide phenylhydrazone (FCCP) and 0.5μM/0.5μM rotenone/antimycin A (all XFp Cell Mito Stress Test Kit, Agilent Technologies, 103010-100) according to manufacturer’s instructions.

#### Glycolytic Rate

Glycolysis-derived proton efflux rate (PER) was determined in primary astrocytes by measuring ECAR under basal conditions and in response to 0.5μM/0.5μM rotenone/antimycin A and 50 mM 2-deoxyglucose (2-DG) (all XFp Glycolytic Rate Assay Kit, Agilent Technologies, 103346-100) according to manufacturer’s instructions. ECARs were converted into PERs and GlycoPERs (glycolysis-specific PERs) by taking the buffer capacity of the medium into account and using the Seahorse XFp Glycolytic Rate Assay Report Generator.

#### ATP Production Rate

Mitochondrial respiration-derived and glycolysis-derived ATP production rates were determined in primary astrocytes by measuring OCR and ECAR under basal conditions and in response to 1.5 μg/mL oligomycin and 0.5μM/0.5μM rotenone/antimycin A (all XFp Real-Time ATP Rate Assay Kit, Agilent Technologies, 103591-100) according to manufacturer’s instructions. OCRs and ECARs were converted to ATP production rates using the Seahorse XFp Real-Time ATP Rate Assay Kit Report Generator.

#### Fuel Dependency

Fuel dependency indicates at what extent cells are unable to compensate for a blocked fuel oxidation pathway using the other non-blocked fuel oxidation pathways. The rate of fatty acids and glutamine oxidation dependency by primary astrocytes was determined by measuring OCR under basal conditions and in response to a different order of combinations of 2 μM UK-5099 (Sigma-Aldrich, PZ0160), 4 μM etomoxir (Sigma-Aldrich, E1905) and 3 μM BPTES (Sigma-Aldrich, SML0601) (all XFp Mito Fuel Flex Test Kit, Agilent Technologies 103270-100) according to the manufacturer’s instructions. Etomoxir was applied first in fatty acid dependency assay and BPTES in glutamine dependency and a combination of the two remaining compounds was applied afterwards.

#### Glutamine Capacity

Glutamine capacity indicates at what extent cells are able to oxidize glutamine when the other fuel (glucose and fatty acids) oxidation pathways are blocked. The rate of glutamine oxidation capacity by primary astrocytes was determined by measuring OCR under basal conditions and in response to a first injection containing 2μM/4μM UK-5099/etomoxir, followed by another containing 3 μM BPTES (all XFp Mito Fuel Flex Test Kit, Agilent Technologies, 103270-100) according to manufacturer’s instructions.

### L-Lactate Release

L-lactate release was assessed in primary astrocyte cultures following two different methods: Firstly, astrocytes were exposed to fresh complete culture medium, and medium was collected after 2, 4 or 7 consecutive days of exposure. L-lactate accumulated in the medium was detected using a Cobas c311 Analyser (Roche). Secondly, a separate experimental group of astrocytes was incubated in DMEM without glucose (GIBCO, 11966025) for 6 h, and then exposed to 4.5 mM glucose overnight. The following morning, 20 μL of supernatant per sample were subjected to the colorimetric EnzyChrom^TM^ Lactate Assay Kit (BioAssay Systems ECLC-100) following the manufacturer’s instructions.

### 2-DeoxyGlucose (2-DG) Uptake

Glucose uptake by cultured primary astrocytes was studied using a bioluminescent method with the Glucose Uptake Assay Kit (Promega, J1341) according to manufacturer’s instructions. Briefly, during the 6 h that preceded the assay, astrocytes were incubated in DMEM without glucose (GIBCO). Thereafter, astrocytes were incubated with 2-deoxyglucose (2-DG), whose uptake and rapid metabolism leads to 2-deoxyglucose-6-phosphate (2-DG6P) accumulation. Afterwards, 2-DG6P was oxidized leading to NADPH generation, and NADPH was determined by an enzymatic recycling amplification reaction using a luminometer.

### ATP Determination

To assess the ability of primary astrocytes to release ATP, primary astrocytes cultured in 12-well plates were washed with HBSS (GIBCO, Thermo Fisher Scientific, 14170-088) and subjected to starvation in HBSS for 4 hours. Afterwards, the medium was replaced with HBSS containing 6-N,N-Diethyl-β-γ-dibromomethylene-D-adenosine-5′-triphosphate (ARL 67156; Tocris, 1283) at a concentration of 100 μM, and either vehicle, 1 μM insulin (Actrapid®, Novo Nordisk) or 1 μM insulin + 10 nM S961 (Phoenix Pharmaceuticals, 051-86). After 30 min of incubation, supernatant was collected and ATP content was quantified using the luciferase-based ATP Determination Kit (Invitrogen, A22066) following the manufacturer’s instructions.

To assess ATP content in *in vivo* microdialysed samples, samples were thawed at RT and immediately subjected to the same ATP Determination Kit following manufacturer’s instructions.

### ^18^F-FDG PET

2-Deoxy-2-[^18^F]fluoro-D-glucose (^18^F-FDG) and positron emission tomography (PET) were used to assess brain glucose metabolism. Briefly, mice were fasted overnight, and the following morning ^18^F-FDG (9,5 MBq ± 0,6 in 80–100 μL) was injected intravenously. After an uptake period of 50 min, mice were anesthetized with 2% isoflurane in 100% O_2_ gas and placed in a small PET tomograph (Mosaic, Philips). Mice were subsequently subjected to a 15 min PET acquisition. Images were reconstructed applying dead time, decay, and random and scattering corrections. Images were semi-quantitatively analysed using the PMOD v3.2 software (PMOD Technologies Ltd), expressing values in standardized uptake value (SUV) units, using the formula SUV = [tissue activity concentration (Bq/cm^3^)/injected dose (Bq)] × body weight (g). ^18^F-FDG metabolism was assessed by drawing two spherical volumes of interest (VOIs) for each image including the entire brain and the reference organ (liver). Semiautomatic delineation was applied with a threshold of 50% of the maximum or minimum voxel value to obtain new VOIs that delimited brain or the reference organ, respectively. The average SUV (SUV_mean_) of the voxels within the VOIs was calculated, and SUV_mean_ ratios were calculated by dividing brain SUV_mean_ by reference organ SUV_mean_. After PET acquisition, mice were euthanized and their brains were dissected to assess *ex vivo* radiotracer incorporation across brain areas (hypothalamus, hippocampus, frontal cortex, and cerebellum). A gamma counter (Hidex Automatic Gamma Counter, Hidex Oy) was used to perform *ex vivo* counting of radioactivity in the samples, which were dissected by the same experimenter and corrected by sample weight. *Ex vivo* radiotracer incorporation was expressed as percentage of the injected dose.

### Assessment of Body Composition

Lean and fat mass body composition were analysed using the EchoMRI^TM^-100H Analyser (EchoMRI) according to manufacturer’s instructions. Mice were weighted and carefully placed into the compact Analyser chamber. Mice weight was introduced to the MRI analysis program and the MRI scanning was immediately started, lasting for 2.5-3 minutes and measuring both lean and fat masses in grams. All analyses were performed in 17- to 18-week-old mice at the same daytime.

### Fasting-Induced Hyperphagic Response

Feeding behaviour was assessed with fasting-refeeding experiments. To this end, the afternoon before the test day, bedding material was changed and mice were fasted overnight for 12 h. At about 8:00 a.m., food was added and its consumption was measured for the following 4 h.

To assess glucose suppression of fasting-induced hyperphagia, mice were fasted overnight for 12 h. The following day, mice were injected with either vehicle (0.9% saline) or glucose (2g/kg body weight) i.p. at 8:00 a.m., and food was provided 30 min after. Subsequently, food was weighted every hour during 4 h in order to measure consumption.

In all cases, food intake was expressed as kcal (normal chow diet = 2.7 kcal/g; high fat diet = 5.21 kcal/g) consumption corrected per animal body weight in grams (kcal/g BW).

### Glucose and Insulin Metabolic Tests

#### Glucose Tolerance Test (GTT)

16- to 17-week-old mice were subjected to a fasting period of 6 h. After fasting period, a subtle puncture in the tail of each animal was performed, and a drop of tail blood was collected with a handheld glucometer (Aviva; Accu-Chek) and blood glucose test strips (Roche, 06453970037) to measure fasting glucose levels (mg/dL) (time 0). Then, mice received an intraperitoneal (i.p.) injection of D-glucose in 0.9% saline at a dose of 2g/kg body weight. Blood glucose levels were measured by tail puncture at the following time points: 15, 30, 60 and 120 min after injection.

#### Insulin Tolerance Test (ITT)

16- to 17-week-old mice were subjected to a fasting period of 1 hour, after which a subtle puncture in the tail of each animal was performed, and a drop of tail blood was collected with a handheld glucometer (Accu-Chek) to measure glucose levels (time 0). Then, they were i.p. injected with insulin (Actrapid®, Novo Nordisk) in 0.9% saline at a dose of 0.75U/kg body weight. Glucose levels were measured at the following time points: 15, 30 and 60 min after injection.

#### Glucose-Stimulated Insulin Secretion (GSIS)

16- to 17-week-old mice were subjected to a fasting period of 6 hours, after which a subtle puncture in the tail of each animal was performed, and a blood sample (minimum quantity: 12 μL of blood) was collected (time 0). Then, mice received an intravenous (i.v.) injection of D-glucose in 0.9% saline at a dose of 1g/kg body weight. Blood samples (12 μL) were collected at the following time points: 2, 5, 15 and 30 min after injection. Blood glucose levels were measured at each time point using a handheld glucometer (Accu-Chek) to ensure that glucose levels first increased after injection and subsequently decreased. Mice that failed to increase blood glucose levels during the test were discarded. Blood samples were immediately centrifuged (10 min, 3500 rpm) and the resulting supernatant (serum) was collected and stored at -80°C until determination of insulin levels.

### Serum and Cerebrospinal Fluid Analyses

#### Serum Insulin Levels

Serum was collected as described above (see “Glucose-Stimulated Insulin Secretion (GSIS)”). Insulin levels were assessed by enzyme-linked immunosorbent assay (ELISA) using the Mouse Ultra-Sensitive Insulin ELISA (Crystal Chem, 90080) according to manufacturer’s instructions.

#### Cerebrospinal Fluid Glucose Levels

18-week-old mice received an i.p. injection containing either vehicle (0.9% saline) or glucose (2g/kg body weight). After 30 minutes, mice were anesthetized using i.p. ketamine/xylazine 80/5 mg/kg respectively, and placed in a stereotaxical apparatus (David Kopf Instruments) to facilitate micro-dissection. Mice were carefully micro-dissected to expose the dura above the cisterna magna. Cerebrospinal fluid (CSF) was collected from the cisterna magna by using a glass micro-capillary. Once collected, samples were microscopically inspected to detect and discard blood-contaminated samples. Non-contaminated samples were immediately frozen and stored at -80°C until glucose determination. Glucose levels were assessed using the colorimetric Glucose Assay Kit (Abcam, ab65333) according to manufacturer’s instructions.

### Cold Exposure

Core body temperature was measured in 18- to 20-week-old animals using a rectal probe attached to a digital thermometer (Panlab, Harvard Apparatus, TMP812RS). Immediately after the first temperature measurement (−1 h), mice were fasted, and a second measurement was performed one hour later (0 h). Immediately, mice were placed into a 4°C room, in which they were kept for the following 4 h. During this time, temperature was measured hourly.

### Behavioural Tasks

All behavioural experiments were carried out between 8 a.m. and 12 p.m. Researchers were blind to mice genotypes and treatments. All tests were undertaken in the same mice cohort at the age of 18-22 weeks.

#### Open Field

In order to examine locomotor activity, mice were individually placed in a light-toned, wood-made open field (35 x 35 cm, 45 cm height), being allowed to move freely for 30 min. The room was dimly illuminated, and a video-tracking system (EthoVision XT version 11.5, Noldus Information Technology) was used to record all animal activity and to analyse the total distance travelled (in cm) for each mouse during the 30-min period.

#### Novel Object Recognition (NOR)

The NOR task was used to test cognition and particularly to examine recognition memory, as previously described^61, 62^. Animals performed the NOR task on the following day to the Open Field experiment. The NOR task took place in the same wood-made open field described for the Open Field experiments, at the same dimly-illuminated room, and thus animals were already familiarized with the open field. The day of the NOR task, animals were subjected to a 5 min sample phase in which they were allowed to freely explore two identical objects placed at identical positions with respect to the open field walls. 1 h later, one of the objects was replaced by another novel, unfamiliar object, and mice were again allowed to freely explore the object-containing open field for 5 min. 24 h later, the replaced object was replaced by another different, unfamiliar object (and both objects interchanged their position one with the other). Mice were then allowed to freely explore the object-containing open field for 5 min. All trials were recorded using a video-tracking system (EthoVision XT version 11.5, Noldus Information Technology). The exploration time that mice dedicated to an object was defined by the time during which mice oriented their snout toward the object at a distance less or equal to 2 cm. Results were expressed as discrimination index, calculated by dividing the time dedicated to explore the novel object by the total object exploration time, and multiplying the result 100 times to obtain a percentage.

#### Morris Water Maze (MWM)

The Morris Water Maze was used to test spatial memory and to evaluate the working and reference memory functions as previously described ^50^. The water maze consists of a 100-cm diameter circular pool (Ugo Basile SRL, 40105) filled with water (20°C). To ensure that the animals owned proper vision and swimming abilities, mice were first subjected to a visible-platform training (habituation phase) conducted with the platform placed in the north-east quadrant over 6 consecutive trials. No visual cues were placed on the walls surrounding the pool. Each trial was terminated when the mouse reached the platform (escape latency) or after 60 s. Mice failing to reach the platform were guided onto it. After each trial, mice remained on the platform for 10 s. On the following days, hidden-platform training (acquisition phase) was conducted over 9 consecutive days (4 trials/day). Trials were conducted following the same indications described above, but with the platform placed in the south-west quadrant 1 cm below the water surface, and with two large visual cues placed on the east and west walls surrounding the pool to guide the mice to the hidden platform. To test memory retention, retention trials were conducted at days 4, 7 and 10 (probes 1, 2 and 3, respectively). In the retention trial, the platform was removed from the pool and mice were allowed to swim freely for 60 s, and the percentage of time spent in the south-west quadrant (target quadrant) was monitored. All trials were recorded using a video camera above the centre of the pool and connected to a video tracking system (EthoVision XT version 11.5, Noldus Information Technology).

### Adeno-Associated Virus (AAV) Injection

AAV injection in the arcuate nucleus of the hypothalamus (ARC) was performed as previously described ^25^. Briefly, 8-week-old GLUT1^f/f^ mice were anesthetized with ketamine/xylazine 80/5 mg/kg i.p. and were placed into a small stereotaxic apparatus (David Kopf Instruments). 1 μL of either AAV5-GFAP-Cre-IRES-GFP or AAV5-GFAP-GFP (custom preparations generated at Viral Vector Production Unit, Universitat Autònoma de Barcelona) was injected bilaterally into the ARC using the following coordinates: anteroposterior (AP): -1.5 mm; mediolateral (ML): +/-0.3 mm; dorsoventral (DV): -5.7 mm. AAV5-GFAP-GFP-injected GLUT1^f/f^ mice were used as a control group.

### Intracerebroventricular (i.c.v.) Injections

#### Cannulation

16- to 17-week-old mice were subjected to brain cannulation (intra-lateral ventricular cannulation). To this end, animals were anesthetized with ketamine/xylazine 80/5 mg/kg i.p. and placed into a stereotaxic apparatus (David Kopf Instruments). 26-gauge guide cannulae (Plastics One, C315GS-5/SPC) were implanted to the left lateral ventricle using the following coordinates: anteroposterior (AP): -0.4 mm; mediolateral (ML): +1.0 mm; dorsoventral (DV): - 2 mm. Immediately, dental acrylic was applied to adhere the guide cannula to the skull surface. Once the dental acrylic dried, a dummy cannula (Plastics One, C315DC/SPC) was inserted into the guide cannula to prevent entry of material into the guide cannula when not in use. Animals were allowed to recover for a minimum of 7 days prior to being subjected to any experiment. Stereotaxic implants were anatomically verified after experiments were completed.

#### Drug Administration

For i.c.v. injections, a 33-gauge Internal Injector (Plastics One, C315IS-5/SPC) was inserted into the guide cannula and a Hamilton Syringe was used to administrate 1 μL of 2 mM pyridoxalphosphate-6-azophenyl-2’,4’-disulfonic acid (PPADS, Tocris, 0625), 1.5 μg/μL S961 (Phoenix Pharmaceuticals, 051-86), or 20 μM 2-methylthyo-adenosine-5’-triphosphate (2-MeSATP, Tocris, 1062) over a 5-min period. Injections were performed 30 min before a behavioural experiment started, 30 min before saline or glucose administration (i.e. 1 hour ahead of refeeding) in fasting-refeeding experiment or 30 min before the baseline (time = 0) blood glucose measurement in GTT and ITT experiments. For both GTT and ITT, in order to test if administered drugs could affect blood glucose levels, an extra baseline blood glucose measurement was made immediately prior to i.c.v. drug injection (time = -30 min).

### *In Vivo* Microdialysis

18-week-old mice were subjected to *in vivo* microdialysis. To this end, animals were anesthetized with ketamine/xylazine 80/5 mg/kg i.p. and placed into a stereotaxic apparatus (David Kopf Instruments). An intra-cerebral guide cannula (Microbiotech/se AB, MAB 4.15.IC) was implanted to the frontal cortex using the following coordinates: anteroposterior (AP): +2.0 mm; mediolateral (ML): +0.3 mm; dorsoventral (DV): -3.3 mm. Immediately, dental acrylic was applied to adhere the guide cannula to the skull surface. Cannulae included a dummy cannula to prevent entry of material into the guide cannula when not in use. After a minimum of 7 days of recovery, mice were placed in the microdialiysis mouse cage and fasted overnight for 12 h. The following morning, a dialysis probe with a cuprophane membrane (Microbiotech/se AB, MAB 4.15.2.Cu) was inserted through the guide cannula, and animals were subjected to a stabilization period by being perfused with artificial cerebrospinal fluid (aCSF. In mM: 148 NaCl, 2.7 KCl, 1.2 CaCl_2_ and 0.85 MgCl_2_. pH = 7.4 adjusted with 1 mM K_2_HPO_4_) for 2 h at a rate of 0.5 μL/min using a glass Hamilton syringe connected to an infusion pump. To abolish ATP metabolism, an ecto-ATPase inhibitor, 6-N,N-Diethyl-β-γ-dibromomethylene-D-adenosine-5′-triphosphate (ARL 67156; Tocris, 1283) was perfused through the probe at concentrations of 1 mM. After the 2 h stabilization period, samples started being collected every 30 min during 1 h through a tube connected to the probe in order to measure basal ATP levels. After 1 h of sample collection, mice were injected glucose i.p. (2g/kg body weight) dissolved in 0.9% saline, and samples were collected every 30 min for another 1 h period. Samples were stored at -80°C until ATP content determination. Mice were conscious and freely moving during the whole sample collection period. Stereotaxic implants were anatomically verified after experiments were completed.

### Cell Sorting

#### Fluorescence-Activated Cell Sorting (FACS)

To isolate astrocytes from brains, a modified version of previously described protocols^63, 64^ was followed. First, anesthetized mice were transcardially perfused with ice-cold PBS for 30 min. Following, brains were extracted and dissected to obtain the corresponding hippocampus, frontal cortex and hypothalamus, which were immediately chopped into tiny fragments in a Petri plaque containing a homogenization solution (in mM: 116 NaCl, 5.4 KCl, 26 NaHCO_3_, 1 NaH_2_PO_4_, 1.5 CaCl_2_, 1 MgSO_4_, 0.5 EDTA, 25 glucose, 1 L-cysteine) with papain (20 U/ml, Sigma-Aldrich, P3125) and DNase I (150 U/μl; Roche, 10104159001) for digestion at 37°C for 15 min. This process was helped by careful pipetting. Then, tissue clogs were removed by filtering the sample through a 70 μm cup Filcon (BD Biosciences) to a 15 mL Falcon tube with 5 mL of 20% FBS in HBSS to quench the reaction. Further astrocyte enrichment was accomplished by removing myelin with Percoll^TM^ (GE Healthcare, GE17-0891-02) gradients. Samples were centrifuged using a swinging bucket for 5 min at 200 x *g* and cell pellet was resuspended in a 20% solution of isotonic Percoll (20% SIP; in HBSS), obtained from a previous stock of SIP (9 parts of Percoll per 1 part of PBS 10X (GIBCO, 14200-067)). Immediately afterwards, samples were smoothly and slowly layered with HBSS poured using Pasteur pipettes. Then, these gradients were centrifuged for 20 min at 200 x *g* with minimum acceleration and no brake, to prevent interphase disruption. The resulting interphase was discarded, and cells were washed in HBSS and centrifuged for 5 min at 200 x *g.* Afterwards, pellet was resuspended in 100 μl of PBS containing the appropriate antibodies: anti-ACSA-2-APC (1:50, Miltenyi Biotec, 130-117-535) and anti-GLUT1 conjugated to Alexa Fluor 488^TM^ (1:200, Abcam, ab195359). Samples were incubated with antibodies for 30 min at 4°C in the dark. Then, PBS was added to further dilute antibodies and stop antibody binding, and samples were centrifuged 10 min at 1500 rpm. Pellets were finally resuspended in sorting buffer (25 mM HEPES, 5 mM EDTA and 1% BSA in HBSS) and 2μL of 7-aminoactinomycin D (7-AAD, Thermo Fisher Scientific, A1310) at a concentration of 0.1 mg/mL were added. Astrocyte cell sorting was performed using a FACS Aria III (BD Biosciences) in the single-cell mode at the adequate sorting rate. A negative control of non-antibody-incubated cells was used to determine background fluorescence. Astrocytes were isolated following the subsequent sorting strategy: Firstly, forward scatter height (FSC-H) was plotted against forward scatter area (FSC-A) to select the appropriate population to sort single cells from cell clusters or debris. Secondly, the adequate subpopulation was sorted based on cellular complexity by plotting side scatter area (SSC-A) against FSC-A. Within the selected subpopulation, dead cells were identified as positive for 7-AAD fluorescence signal and were discarded. Finally, cells with high (>10^3^) ACSA-2-APC fluorescence signal were identified as astrocytes and were sorted. Sorted cells were collected either in Lysis Buffer (RNeasy^®^ Micro Kit, QIAGEN 74004) or PBS. Cells collected in Lysis Buffer were stored at -80°C until RNA extraction. Cells collected in PBS were immediately centrifuged 2 min at 13000 rpm to generate a pellet, supernatant was discarded and the dry pellet was stored at -80°C until protein extraction. Posterior analyses using FlowJo 10.8.1 were performed to identify ACSA-2-APC-positive cells with high (>10^3^) GLUT1-AlexaFluor^TM^ 488 fluorescence signal as GLUT1-positive astrocytes.

#### Magnetic-Activated Cell Sorting (MACS)

To isolate astrocytes from brains ensuring a gentle and not reactivity-triggering process for an accurate astrocyte reactivity gene profile analysis, MACS was performed. First, anesthetized mice were transcardially perfused with ice-cold PBS for 30 min. Following, brains were extracted and dissected to obtain the corresponding hippocampus, frontal cortex and hypothalamus, which were immediately chopped and digested at 37°C with rotation in a solution containing 2mg/ml papain (Worthington Biochemical Corporation, LS003126) and 50 μg/ml DNase I in PBS. Then, the tissue was mechanically disaggregated and tissue clogs were removed by filtering the sample through a 70 μm cup Filcon (BD Biosciences). Cell suspensions were centrifuged 15 min at 300 x *g*. The resulting cell pellets were resuspended in 25% Percoll and centrifuged 10 min at 1000 x *g*. After cell debris and myelin removal, cell pellets were resuspended in sorting buffer and incubated with ACSA-2 MicroBeads (1:10, Miltenyi Biotec, 130-097-678) for 15 min at 4°C. Following, astrocyte ACSA-2^+^ cell sorting was performed using an autoMACS Pro Separator (Miltenyi Biotec). After isolation, cells were pelleted and resuspended in the Homogenization Solution of Maxwell^®^ RSC simplyRNA Tissue Kit (Promega, AS1340) and stored at -80°C until quantitative PCR experiments.

### Quantitative PCR (qPCR)

RNA extraction from cell samples was performed using the RNeasy^®^ Micro Kit (Qiagen) or Maxwell^®^ RSC simplyRNA Tissue Kit (Promega, AS1340) according to manufacturer’s instructions. RNA extraction from brown adipose tissue (BAT) samples was performed using the TRI Reagent^®^ (Sigma Aldrich, T9424). Subsequently, RNA samples were subjected to retro-transcription using the High-Capacity cDNA Reverse Transcription Kit (Applied Biosystems, Thermo Fisher Scientific, 4368813). Quantitative real-time PCR was performed on CFX384 Touch Real-Time PCR Detection System (Bio-Rad). TaqMan^®^ Universal PCR Master Mix (Applied Biosystems, 10733457) was used with TaqMan^®^ Probes for *Gapdh* (Applied Biosystems, Mm99999915_g1), *Insr* (Applied Biosystems, Mm01211875_m1), *Slc2a1* (Applied Biosystems, Mm00441480_m1), *Slc2a2* (Applied Biosystems, Mm00446229_m1), *Slc2a3* (Applied Biosystems, Mm00441483_m1), *Slc2a5* (Applied Biosystems, Mm00600311_m1) and *Ucp1* (Applied Biosystems, Mm01244861_m1). iQTM SYBR^®^ Green Supermix Reagent (Bio-Rad, 1708880) was used with the SYBR probes detailed in Table S1. Sequences for the probes used to assess mRNA expression of the different transporters of the GLUT family were obtained from^65^. Sequences for the probes used to assess mRNA expression of astrocyte reactivity genes were obtained from^66^.

Relative quantification of the gene expression of all targets with respect to the control group was performed by a comparative method (2^-ΔΔCt^). *Gapdh* was used as an internal control for all experiments except for the experiment assessing the mRNA expression of astrocyte reactivity genes where *Actb* was used as internal control.

### Brain Capillary Depletion

To assess protein specifically in brain capillaries and in capillary-depleted brains, brain capillaries were separated using the previously described brain capillary depletion technique^67–69^ with minor modifications. Briefly, one hemisphere per mouse was mechanically homogenized in 3.5-fold excess volume of ice-cold homogenization buffer (10 mM HEPES, 141 mM NaCl, 4 mM KCl, 2.8 mM CaCl_2_, 1 mM MgSO_4_, 1 mM NaH_2_PO_4_, and 10 mM glucose in dH_2_O; pH 7.4) in a glass dounce tissue grinder. After homogenization, ice-cold dextran (MW: 60000, Sigma-Aldrich, 31397) was added in an equal volume in a concentration of 26%, and the mixture was further homogenized. Next, homogenates were centrifuged at 5800 x *g* for 15 min at 4°C, providing slow deceleration. The capillary-free brain homogenate supernatant was collected separately from the capillary-containing pellet. The pellet was washed three times with 1.5 mL ice-cold buffered solution. The capillary-free supernatant was resuspended in 10 mL ice-cold buffered solution and centrifuged 1,500 x *g* in order to obtain the cells. The capillary-containing and capillary -depleted samples were then resuspended in ice-cold buffered solution, sonicated and centrifuged at 20000 x *g* for 20 min. Supernatant coming from both types of samples were stored at -80°C until protein content analysis and subsequent Western blotting.

### Western Blotting

Protein extraction was performed by homogenization of cells or tissue in ice-cold RIPA buffer (50 mM Tris-HCl pH = 7.4; 0.25% deoxycholate; 1% Nonidet P-40; 150 mM NaCl; 1 mM EDTA, 1 mM PMSF; 1 μg/mL aprotinin, 1 mM Na_3_VO_4_, 1 mM NaF), followed by a 30-min incubation on ice. Then, samples were centrifuged (14000 x *g*, 20 min) at 4°C, and the resulting supernatant was collected and stored at -80°. Sample protein content was measured using the Bio-Rad Protein Assay (Bio-Rad, 5000006) according to manufacturer’s instructions. Equal amounts of protein (15 μg per sample) were separated by electrophoresis using sodium dodecyl sulphate – polyacrylamide (7,5%) gels under reducing conditions, and transferred onto Amersham^TM^ Protran^TM^ Premium 0.45 μm nitrocellulose membranes (GE Healthcare, GE10600048) for 16 h at 30V. The resulting trans-blots were blocked in a 5% powder milk-containing TBS-Tween 20 (TBS-T) solution for 1h to prevent antibodies from non-specific binding to the membrane. Membranes were incubated overnight at 4°C with the corresponding primary antibody. The following primary antibodies and dilutions were used: anti-GLUT1 (1:1000, Merck Millipore, 07-1401), anti-UCP1 (1:1000, Abcam, ab209483), anti-fibrinogen (1:1000, Dako, Agilent Technologies, A0080), anti-ZO-1 (1:1000, Thermo Fisher Scientific, 40-2200), anti-Occludin (1:1000, Thermo Fisher Scientific, 40-4700). The following day, membranes were washed in TBS-T, and subsequently incubated 2h at RT with secondary antibodies conjugated to IRDye 800CW or IRDye 680CW (LI-COR Biosciences, Lincoln, NE, USA) diluted to 1:5000 in TBS with 5% BSA. Bands were visualized using Odyssey^®^ DLx Imaging System (LI-COR Biosciences), and Optical Density (O.D.) was analysed using Image Studio^TM^ Lite (LI-COR Biosciences). β-actin (1:5000, Sigma-Aldrich, A5441) was used as internal loading control.

Original raw blots have been deposited at Mendeley (DOI: 10.17632/zdc9tmktjp.1) and can also be found in Source data.

### Histology

#### Primary Astrocyte Immunofluorescence

5000 astrocytes/well were seeded in 12-well removable chamber slides (Ibidi 81201) pre-coated with 50 μg/mL poly-D-lysine and incubated for 48 h in complete culture medium at 37°C and 5% CO_2_. Thereafter, medium was removed, cells were washed in PBS and fixed using 4% PFA for 20 min at RT. Next, cells were washed in PBS and incubated with blocking buffer (1% BSA and 0.1% Triton-X-100 in PBS) at RT for 1 h. Following, cells were incubated overnight at 4°C with primary antibodies diluted in blocking buffer at the appropriate concentration. The following primary antibodies and dilutions were used: anti-GFAP (1:1000, Cell Signalling Technology, 3670) and anti-GFP (1:500, Molecular Probes, A-11122). The next day, cells were then washed in PBS and incubated with secondary antibodies for 60 min at RT under light-protected conditions. The following secondary antibodies and dilutions were used: Donkey anti-Mouse IgG conjugated to Alexa Fluor^TM^ 546 (1:200, Invitrogen, Thermo Fisher Scientific, A10036) and Donkey anti-Rabbit conjugated to Alexa Fluor^TM^ 488 (1:200, Invitrogen, Thermo Fisher Scientific, A21206). After a final washing 3 x with PBS for 10 min, cells were mounted with DAPI Fluoromount-G^®^ Mounting Media (Southern Biotech, 0100-20). All images were acquired using a confocal laser scanning microscope (LSM 800 with Airyscan, Zeiss) equipped with a 40x objective, and the ZEN 2 Blue Edition software (Zeiss). To ensure similar imaging conditions for all images of all slides, an identical microscope set-up (including objective, zoom, laser power and gain) was used to acquire all pictures within the same experiment. All images were acquired using a pinhole of 1 airy unit. In order to obtain representative images, maximum intensity projections were made in FIJI^70^ and identically adjusted for brightness and contrast.

#### Pancreas and Brown Adipose Tissue Immunohistochemistry

6 hour-fasted mice were euthanized, and both brown adipose tissue and pancreas were carefully dissected. Brown adipose tissue and pancreas were formalin-fixed, paraffin-embedded and cut in 3-μm thick sections. Brown fat sections were stained with haematoxylin and eosin (H&E) for routine histology. In addition, immunohistochemistry was applied using the following primary antibodies: rabbit monoclonal anti-UCP1 (1:4000, Abcam, ab209483) and rabbit monoclonal anti-insulin (1:80000, Abcam, ab181547). Antigen retrieval was performed in a Pascal pressure chamber (Dako S2800) for 30 min at 95 °C in 10 mM Tris-1 mM EDTA solution with pH = 9 (for UCP1) or in 10 mM citrate acid solution with pH = 6 (for insulin). Slides were allowed to cool for 20 min, then endogenous peroxidase was blocked with 3% H_2_O_2_ in deionized water for 12 min. After washing in TBS-T, sections were incubated overnight at 4°C with primary antibodies. The following morning, sections received a washing in TBS-T, and were subsequently incubated with goat anti-rabbit labelled polymer EnVision+ Single Reagent (Dako, K400311-2) for 30 min at RT and peroxidase activity was revealed using DAB+ (Dako, K346811-2). Finally, sections were lightly counterstained with Harris haematoxylin, dehydrated, and coverslipped with Eukitt (Labolan, 28500).

#### Brain Slice Immunofluorescence

Mice were anesthetized with ketamine/xylazine 80/5 mg/kg i.p. and transcardially perfused with ice-cold PBS followed by 4% PFA in PBS. Brains were extracted and post-fixed overnight in 4% PFA at 4°C, and then were exposed to 30% sucrose in PBS. Brains were sectioned to a thickness of 40 μm using a sliding freezing microtome (Leica SM2010 R) and preserved in cryoprotectant solution (25% glycerol and 30% ethylene glycol in PBS). Free-floating brain slices were washed in PBS. For c-Fos staining, slices were then incubated in 10 mM sodium citrate (pH = 8) in PBS at 60°C for 30 min and washed in PBS. Thereafter, all slices were incubated for 2 h in blocking buffer (1% BSA and 0.1% Triton-X-100 in PBS) at RT, and incubated overnight at 4°C with primary antibodies diluted in blocking buffer. The following primary antibodies and dilutions were used: anti-POMC (1:1000, Phoenix Pharmaceuticals, Inc., H-029-30), anti-Tyrosine Hydroxylase (1:1000, Abcam, ab112), anti-c-Fos (1:500, Synaptic Systems, 226004), anti-GFAP (1:1000, Cell Signalling Technology, 3670), anti-GFP (1:1000, Molecular Probes, A-11122), anti-Fibrin (1:1000, Dako, Agilent Technologies, A0080), anti-ZO-1 (1:200, Thermo Fisher Scientific, 40-2200) and anti-Occludin (1:200, Thermo Fisher Scientific, 40-4700). Slices were then washed in PBS and incubated for 2 h at RT with secondary antibodies diluted in 1% BSA in PBS under light-protected conditions. *Lycopersinon esculentum* Lectin conjugated with Fluorescein (1:200, Vector Laboratories, FL-1171) was incubated together with the secondary antibody. The following secondary antibodies and dilutions were used: Donkey anti-Mouse IgG conjugated to Alexa Fluor^TM^ 488 (1:200, Invitrogen, A21202), Donkey anti-Mouse IgG conjugated to Alexa Fluor^TM^ 546 (1:200, Invitrogen, A10036), Donkey anti-Rabbit conjugated to Alexa Fluor^TM^ 488 (1:200, Invitrogen, A21206), Donkey anti-Rabbit conjugated to Alexa Fluor^TM^ 555 (1:200, Invitrogen, A31572) and/or Goat anti-Guinea Pig IgG conjugated to Alexa Fluor^TM^ 488 (1:1000, Invitrogen, A11073). Finally, after a final washing in PBS for 10 min, cells were mounted with DAPI Fluoromount-G^®^ Mounting Media. All images were acquired using a confocal laser scanning microscope (LSM 800 with Airyscan, Zeiss), and the ZEN 2 Blue Edition software (Zeiss). To ensure similar imaging conditions for all images of all slides, an identical microscope set-up (including objective, zoom, laser power and gain) was used to acquire all pictures within the same experiment. All images were acquired using a pinhole of 1 airy unit. In order to obtain representative images, images were randomly taken from two nonadjacent tissue sections per specimen (n = 4-6 per group), maximum intensity projections were made in FIJI^70^ and identically adjusted for brightness and contrast.

### Astrocyte Morphometric Analysis

Imaging of astrocytes located in the stratum radiatum was performed using brain slice immunofluorescence of GFAP-positive cells and confocal laser scanning microscopy (see “Brain Slice Immunofluorescence”) with a 63x objective. Sholl analysis of astrocytic processes was performed semi-automatedly using FIJI, including the plugin “SNT” (https://imagej.net/plugins/snt), as described previously^71^. Briefly, a Z-stack of images was loaded to FIJI, astrocytes were identified by their distinctive GFAP-positive bushy/star-like shape, and the first ten astrocytes per mouse that fulfilled the subsequent criteria were selected for analysis: (i) its GFAP-positive structure contained one single DAPI-positive nucleus, and (ii) its main structure did not include truncated processes. In order to obtain representative images, images were randomly taken from two nonadjacent tissue sections per specimen (n = 6 per group). The SNT plugin was used to trace the GFAP-positive structure defining the centre of the DAPI staining as the starting point, and selecting the distant point within a GFAP-positive process as the ending point. The SNT plugin automatedly connected the two points following the trace of the immunofluorescent GFAP-positive pathway only if such a pathway existed between the points. Number of processes, total length covered by the processes, process thickness, number of intersections (at increasingly distant points from the DAPI-positive centre) and maximum intersection radius were exported. Morphometric analysis was performed by two independent blinded researchers.

### Dendritic Spine Density Analysis

#### Golgi-Cox Staining and Image Acquisition

Mice were euthanized and brains were dissected, shortly rinsed in PBS and immediately incubated in 4% paraformaldehyde (PFA) in PBS at RT for 72 h. Thereafter, the FD Rapid GolgiStain Kit (FD Neurotechnologies, PK401A) was used according to manufacturer’s instructions. A stack of 80-100 focal images (1 μm-thick z stacks) per dendrite (for a total of 30 dendrites per animal) were acquired from randomly chosen hippocampal dentate gyrus from three nonadjacent tissue sections per mouse (n = 4 per group) using a 100x objective-equipped automated microscope (Axio Imager M1, Zeiss).

#### Dendritic Spine Quantification

Dendritic spines were analysed using the Reconstruct software^72^ as previously described^73^. The Reconstruct software enables unbiased classification of spine types. In brief, spines were counted on randomly selected 10 μm-long segments of secondary dendritic branches, and both spine three-dimensional length and width were measured and processed using Reconstruct to classify spines. Spine analysis was performed by two independent blinded experimenters.

### Whole-Brain Clearing and 3D Blood Vessel Analysis

#### Vessel Staining of Cleared Mouse Brain Slices

Mice were transcardially perfused with PBS (pH 7.4) and 10 mL of 4% paraformaldehyde in PBS. Brains were dissected, then post-fixed with 4% PFA overnight. Then, 500-μm-thick brain slices were sectioned using a vibratome (VT1000S, Leica) and maintained in PBS solution at 4 °C for tissue clearing. Sections were cleared following the CUBIC protocol^74^. This is a two-phased method: delipidation (ScaleCUBIC Reagent-1A: urea 25 wt%, Quadrol 25 wt% and Triton X-100 15 wt% and dH2O) and RI matching (ScaleCUBIC Reagent2: urea 25 wt%, sucrose 50 wt%, triethanolamine 10 wt% and dH2O). First, brain slices were incubated with ScaleCUBIC Reagent-1A for 4 days at 37 °C with gentle shaking followed by PBS washing for 16 h at room temperature. Delipidated sections in PBS were blocked with BSA 4 % and incubated with a 50 ug/ml solution of lectin conjugated with fluorescein (Vector Laboratories, FL1171), 24 h at room temperature. Then, sections continued their clearing process in ScaleCUBIC Reagent2. Finally, the slices were immersed in ScaleCUBIC reagent 2 to continue the clearing process until their visualization in the microscope.

#### Two-Photon Excitation Microscopy

Images were collected using a Zeiss LSM 880 (Carl Zeiss, Jena, Germany) equipped with a two-photon femtosecond pulsed laser (MaiTai DeepSee, Spectra-Physics, USA), tuned to a central wavelength of 800 nm using a 25x objective (LD LCI Plan-Apochromat 25x/ 0.8, Carl Zeiss). Tile and z-stack scans from 500 µm sections were acquired in the non-descanned mode after spectral separation and emission re-filtering using 500-550 nm BP filter for Lectin signal.

#### Vessel Image Analysis

Vessel quantification in fluorescent 3D image stacks was carried out using an automatized home-made FIJI plugin. First, 3D images were pre-processed by means of a background subtraction operation followed by a tubeness filter^75^ to enhance tube-like structures. Then, vessels were segmented by thresholding the image using a constant threshold value across all images. The resulting binary mask was further refined by means of a median filter and small particle removal, and the total vessel volume (in μm^3^) and vessel density (as % of tissue occupied by vessels) were calculated. The segmentation mask was finally skeletonized in 3D and the resulting skeleton was analysed^76^ to quantify the number of vessels, the average vessel length (in μm) and the normalized vessel length (in m/mm^3^).

### Proteomic Analysis

#### Protein Extraction

Brain specimens (frontal cortex) derived from control and GLUT1^1′GFAP^ mice were homogenized in lysis buffer containing 7 M urea, 2 M tiourea and 50 mM dithiothreitol (DTT). The homogenates were spinned down at 100.000 x g for 1 h at 15°C. Protein concentration was measured in the supernatants using the Bio-Rad Protein Assay (Bio-Rad, 5000006) according to manufacturer’s instructions.

#### SWATH-Mass Spectrometry Proteomics: MS/MS Library Generation

As an input for generating the SWATH-mass spectrometry (SWATH-MS) assay library, a pool of all samples (10 µg/sample) was generated. Then, the protein extract (30 µg) was diluted in Laemmli sample buffer and loaded into a 0.75 mm thick polyacrylamide gel with a 4% stacking gel casted over a 12.5% resolving gel. Total gel was stained with Coomassie Brilliant Blue and 12 equals slides from the pooled sample was excised from the gel and transferred into 1.5 mL Eppendorf tube. Protein enzymatic cleavage was carried out with trypsin (1:20, w/w; Promega, V5280) at 37 °C for 16 h. Peptide mixture was dried in a speed vacuum for 20 min. Purification and concentration of peptides was performed using C18 Zip Tip Solid Phase Extraction (Millipore). Peptides recovered from in-gel digestion processing were reconstituted into a final concentration of 0.5µg/µL of 2% acetonitrile, 0.5% formic acid, 97.5% MilliQ-water prior to mass spectrometric analysis. MS/MS datasets for spectral library generation were acquired on a Triple TOF 5600+ mass spectrometer (Sciex, Canada) interfaced to an Eksigent NanoLC Ultra 2D Pump System (Sciex, Canada) fitted with a 75 μm ID column (Thermo Scientific 0.075 × 250mm, particle size 3 μm and pore size 100 Å). Prior to separation, the peptides were concentrated on a C18 precolumn (Thermo Scientific 0.1 × 50mm, particle size 5 μm and pore size 100 Å). Mobile phases were 100% water 0.1% formic acid (buffer A) and 100% acetonitrile 0.1% formic acid (buffer B). Column gradient was developed in a gradient from 2% B to 40% B in 120 min. Column was equilibrated in 95% B for 10 min and 2% B for 10 min. During all process, precolumn was in line with column and flow maintained all along the gradient at 300 nl/min. Output of the separation column was directly coupled to nano-electrospray source. MS1 spectra was collected in the range of 350-1250 m/z for 250 ms. The 35 most intense precursors with charge states of 2 to 5 that exceeded 150 counts per second were selected for fragmentation, rolling collision energy was used for fragmentation and MS2 spectra were collected in the range of 230–1500 m/z for 100 ms. The precursor ions were dynamically excluded from reselection for 15 s. MS/MS data acquisition was performed using AnalystTF 1.7 (Sciex) and spectra files were processed through ProteinPilot v5.0 search engine (Sciex) using Paragon^TM^ Algorithm (v.4.0.0.0) for database search. To avoid using the same spectral evidence in more than one protein, the identified proteins were grouped based on MS/MS spectra by the Progroup™ algorithm, regardless of the peptide sequence assigned. The protein within each group that could explain more spectral data with confidence was depicted as the primary protein of the group. False discovery ratio (FDR) was performed using a non-linear fitting method^77^ and displayed results were those reporting a 1% Global FDR or better.

#### SWATH-Mass Spectrometry Proteomics: Quantitative Analysis

Protein extracts (20 µg) from each sample were reduced by addition of DTT to a final concentration of 10mM and incubation at room temperature for 30 minutes. Subsequent alkylation by 30 mM iodoacetamide was performed for 30minutes in the dark. An additional reduction step was performed by 30mM DTT, allowing the reaction to stand at room temperature for 30 min. The mixture was diluted to 0.6M urea using MilliQ-water, and after trypsin addition (Promega) (enzyme:protein, 1:50 w/w), the sample was incubated at 37°C for 16h. Digestion was quenched by acidification with acetic acid. The digestion mixture was dried in a SpeedVac. Purification and concentration of peptides was performed using C18 Zip Tip Solid Phase Extraction (Millipore). The peptides recovered were reconstituted into a final concentration of 0.5µg/µL of 2% acetonitrile, 0.5% formic acid, 97.5% MilliQ-water prior to mass spectrometric analysis. For SWATH-MS-based experiments the instrument (Sciex TripleTOF 5600+) was configured as previously described^78^. Briefly, the mass spectrometer was operated in a looped product ion mode. In this mode, the instrument was specifically tuned to allow a quadrupole resolution of Da/mass selection. The stability of the mass selection was maintained by the operation of the Radio Frequency (RF) and Direct Current (DC) voltages on the isolation quadrupole in an independent manner. Using an isolation width of 16 Da (15 Da of optimal ion transmission efficiency and 1 Da for the window overlap), a set of 37 overlapping windows were constructed covering the mass range 450–1000 Da. In this way, 1 μL of each sample was loaded onto a trap column (Thermo Scientific 0.1 × 50mm, particle size 5 μm and pore size 100 Å) and desalted with 0.1% trifluoroacetic acid at 3 μL/min during 10 min. The peptides were loaded onto an analytical column (Thermo Scientific 0.075 × 250mm, particle size 3 μm and pore size 100 Å) equilibrated in 2% acetonitrile 0.1% formic acid. Peptide elution was carried out with a linear gradient of 2 to 40% B in 120 min (mobile phases A: 100% water + 0.1% formic acid and B: 100% acetonitrile + 0.1% formic acid) at a flow rate of 300 nL/min. Eluted peptides were infused in the mass spectrometer. The Triple-TOF was operated in swath mode, in which a 0.050 s TOF MS scan from 350 to 1250 m/z was performed, followed by 0.080 s product ion scans from 230 to 1800 m/z on the 37 defined windows (3.05 s/cycle). Collision energy was set to optimum energy for a 2 + ion at the centre of each SWATH block with a 15 eV collision energy spread. The mass spectrometer was always operated in high sensitivity mode. The resulting ProteinPilot group file from library generation was loaded into PeakView® (v2.1, Sciex) and peaks from SWATH runs were extracted with a peptide confidence threshold of 99% confidence (Unused Score ≥ 1.3) and a FDR lower than 1%. For this, the MS/MS spectra of the assigned peptides was extracted by ProteinPilot, and only the proteins that fulfilled the following criteria were validated: (1) peptide mass tolerance lower than 10 ppm, (2) 99% of confidence level in peptide identification, and (3) complete b/y ions series found in the MS/MS spectrum. Only proteins quantified with at least two unique peptides were considered. MS data and search results files were deposited in the Proteome Xchange Consortium via the JPOST partner repository (https://repository.jpostdb.org)^79^ with the identifier PXD035376 for ProteomeXchange and JPST001793 for jPOST (for reviewers: https://repository.jpostdb.org/preview/130553713462d6853f72fca; Access key: 1544).

#### Weighted Protein Co-Expression Network Analysis

Weighted gene correlation network analysis (WGCNA) is a robust method developed for the construction of co-expression networks from array-based data. A differential feature of network construction by WGCNA is that it takes into consideration not only the co-expression patterns between two genes but also the overlap of neighbouring genes. We performed WGCNA as previously described^80, 81^. Briefly, given the proteins identified by SWATH-MS, we used the WGCNA package (https://horvath.genetics.ucla.edu/html/CoexpressionNetwork/Rpackages/WGC <u>NA/</u>)^80^ in R (https://www.r-project.org) to build a weighted protein co-expression signed network containing 2329 nodes (proteins). We optimized the parameter “β” to maintain both the scale-free topology of the network as well as sufficient node connectivity, as recommended, selecting β = 8 (scale-free topology R^2^ = 0.904; mean connectivity = 11.8). In such a network, any two proteins were connected and the edge weight was determined by the WGCNA topology overlap measure. These weights ranged from 0 to 1, and considered not only the expression correlation between two connected proteins, but also the amount and connection strength of the “co-neighbour” connected proteins that these two proteins shared. Given a network, we used the Dynamic Tree Cut algorithm to identify modules (clusters) of highly co-expressed proteins, built with the default value of SplitDepth for robust module detection included in the WGCNA package (cutHeight = 200; minModuleSize = 30; deepSplit = 2; mergeCutHeight = 0.25). Each module was identified as a branch cut-off of the dendrogram, and named by a unique colour. Given the 27 modules that we obtained, each module’s eigenprotein was defined as the first principal component of the module, whose value is a representative of the overall expression profile of the proteins within a given module. Therefore, different eigenprotein values of a given module among genotypes imply different expression profiles of the module among these very genotypes. Then, we computed the correlation between the module eigenprotein value and the genotype (control or GLUT1^1′GFAP^), establishing a p-value cut-off of <0.01. Only 1 out of the 27 modules satisfied this condition, a module termed “blue module” by WGCNA. We also used WGCNA to calculate the strength of the contribution of each protein to the overall co-expression pattern of the module (“module membership”) and the significance of each “blue module” protein for the definition of the genotype, and calculated the correlation between both values. Next, we computed the eigenprotein value of each module and compared the values between genotypes. Finally, we performed a Gene Ontology (GO) Biological Processes enrichment analysis using ShinyGO version 0.76 (http://bioinformatics.sdstate.edu/go/)^82^. The 179 proteins that form the “blue module” were included in the analysis, using the 2329 proteins detected as the universe for the GO analysis. The following criteria were applied: pathway criteria exclusion (FDR cut-off) = 0.05; pathway minimum size = 2; pathway maximum size = 2000.

### Quantification and statistical analysis

All values were expressed as mean ± SEM. Statistical analyses were conducted using GraphPad PRISM (version 9.3.1). Statistical details of experiments can be found in the figure legends. Normality was checked with Shapiro-Wilk’s test. Unless otherwise specified in the figure legends, datasets with only two independent groups were analysed for statistical significance using unpaired, two-tailed Student’s t test. Datasets with more than two groups were analysed using one-way ANOVA, followed by Tukey post hoc test. Datasets with two independent factors were analysed using two-way ANOVA, followed by Tukey post hoc test. Two-way repeated measures ANOVA was performed to detect significant interactions between genotype and time, and multiple comparisons were analysed following Tukey’s post hoc tests. All p-values below or equal to 0.05 were considered significant. * p ≤ 0.05, **p ≤ 0.01 and *** p ≤ 0.001.

### Data Availability

Proteomics data have been deposited at ProteomeXchange/jPOST and are publicly available with the identifier PXD035376 for ProteomeXchange and JPST001793 for jPOST (https://repository.jpostdb.org/preview/130553713462d6853f72fca; Access key: 1544).

Original western blot images have been deposited at Mendeley and are publicly available (DOI: 10.17632/zdc9tmktjp.1). Microscopy data reported in this paper will be shared by the lead contact upon request.

### Code Availability

The code used to build a weighted protein co-expression signed network is the WGCNA package

(https://horvath.genetics.ucla.edu/html/CoexpressionNetwork/Rpackages/WGC NA) in R (https://www.r-project.org).

## Supplementary Material

**Fig. S1.**
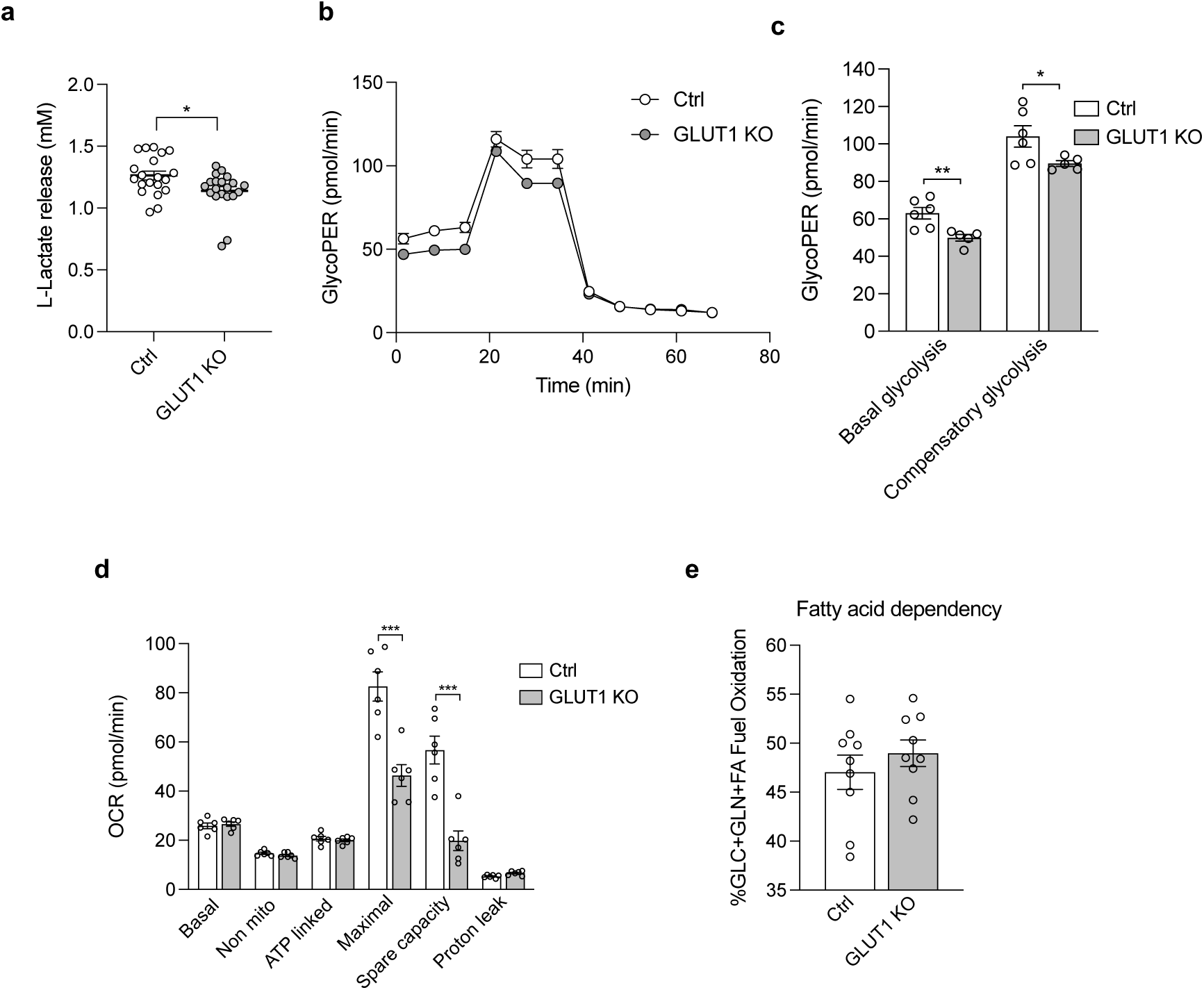
Astrocytic GLUT1 ablation induces a metabolic shift in primary cultured astrocytes. **a**, L-Lactate release in culture medium after glucose starvation and subsequent glucose stimulation (in mM; n = 20 independent wells). **b,** Glycolysis-derived proton efflux rate (GlycoPER) assessment and **c**, quantification of basal and compensatory glycolysis in Ctrl and GLUT1 KO cultured astrocytes (n = 6 independent wells). **d,** Quantification of mitochondrial respiration evaluation (measured by OCR) in Ctrl and GLUT1 KO cultured astrocytes (n = 6 independent wells). **e,** Fatty acid dependency assay (n = 9 independent wells). Data are presented as mean ± SEM. *p ≤ 0.05, ***p<0.001 as determined by two-tailed Student’s t test in all cases.

**Fig. S2.**
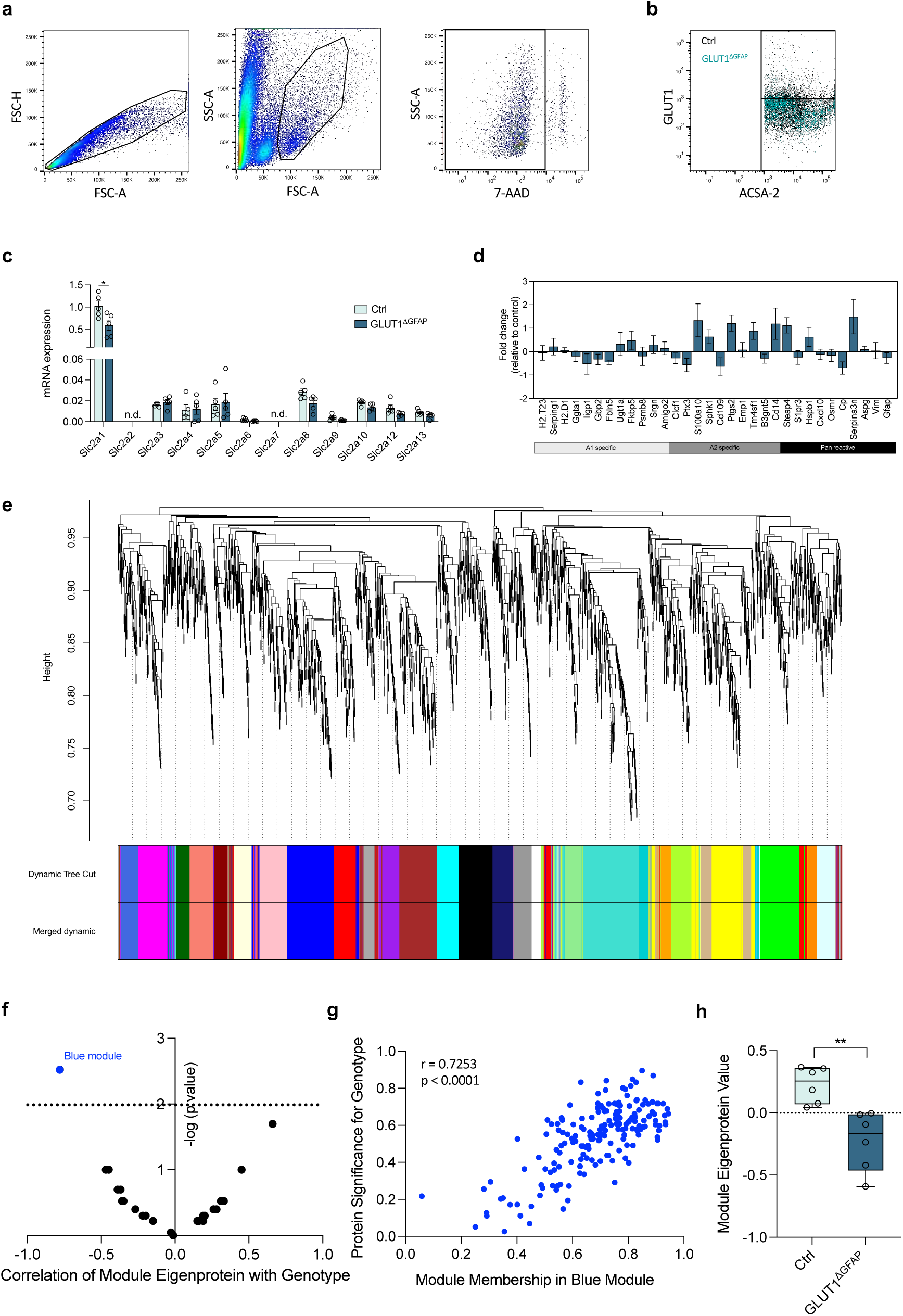
Astrocyte isolation, gene profiling and proteomic analysis. **a**, Astrocyte sorting strategy: Firstly, forward scatter height (FSC-H) was plotted against forward scatter area (FSC-A) to select the appropriate population to sort single cells from cell clusters or debris. Secondly, the adequate subpopulation was sorted based on cellular complexity by plotting side scatter area (SSC-A) against FSC-A. Within the selected subpopulation, dead cells were identified as positive for 7-AAD fluorescence signal and were discarded. Finally, cells with high (>10^3^) ACSA-2-APC fluorescence signal were identified as astrocytes and were sorted. **b,** GLUT1 fluorescence quantification plot in sorted control (Ctrl) and GLUT1^1′GFAP^ astrocytes (n = 6 mice per group). **c,** mRNA expression levels of all GLUT (Slc2a) family members in Ctrl and GLUT1^1′GFAP^ mice (n = 5 mice per group). **d,** Sorted astrocyte gene profiling expressed in fold change of GLUT1^1′GFAP^ compared to control mice (z score = 0) (n = 5 mice per group). **e,** Protein clustering dendrogram built by hierarchical clustering of adjacency-based dissimilarity (WGCNA Dynamic Tree Cut and SplitDepth) to detect 27 co-expression modules (minimum module size = 30 proteins) with corresponding color assignments (n = 6 mice per group)**. f,** Plot representing the r value vs. the p-value of each WGCNA-detected module correlation with the genotype (Ctrl or GLUT1^1′GFAP^) (n = 6 mice per group). **g,** Correlation between protein module membership and protein significance for genotype (Ctrl vs. GLUT1^1′GFAP^ mice) of the 179 proteins composing the blue module (n = 6 mice per group). **h,** Blue module eigenprotein value comparison between Ctrl and GLUT1^1′GFAP^ mice (n = 6 mice per group). Data are presented as mean ± SEM. *p ≤ 0.05, **p ≤ 0.01 as determined by two-tailed Student’s t test (c, d and h). Correlation study was performed using Pearson correlation coefficient (f and g).

**Fig. S3.**
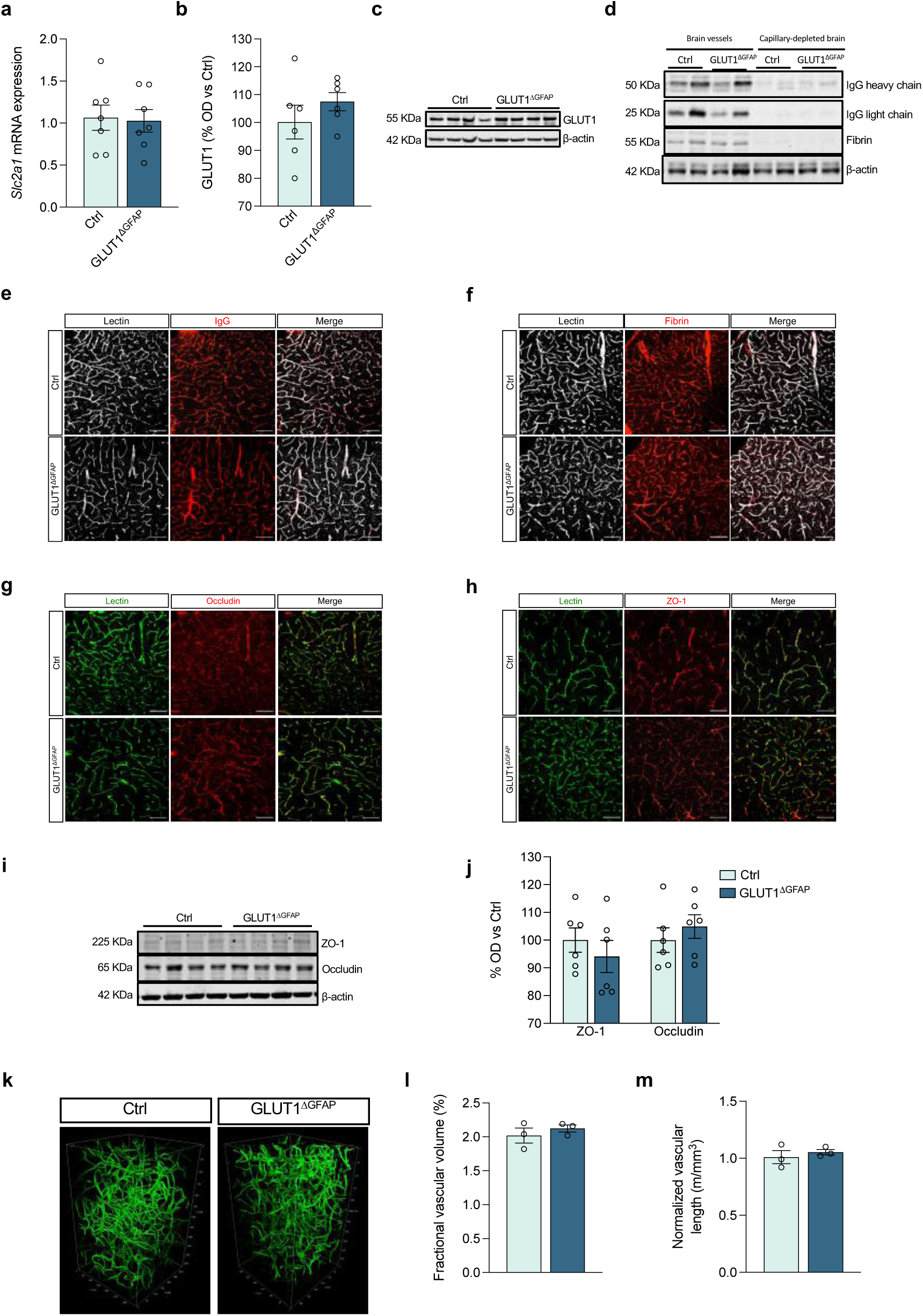
Astrocytic GLUT1 ablation neither induces brain angioarchitecture alteration nor BBB breakdown. **a**, *Slc2a1* mRNA expression levels in isolated brain vessels in control (Ctrl) and GLUT1^1′GFAP^ mice (n = 7 mice per group). **b,** GLUT1 protein expression level quantification (optical density, OD) and **c**, representative western blotting image in Ctrl and GLUT1^1′GFAP^ mice in isolated brain vessels (n = 6 mice per group). **d,** IgG heavy and light chains and fibrin protein expression in homogenized isolated brain blood vessels and capillary-depleted brains of control and Glut1^ΔGFAP^ mice (n = 6 per group). **e,** Lectin(green)-IgG(red) and **f**, lectin(green)-fibrin(red) co-localization in brain sections assessed by immunofluorescence (n = 4 mice per group, two slices per mouse). Scale bar 100 µm. **g,** Lectin(green)-occludin(red) and **h**, lectin(green)-ZO-1(red) co-localization in brain sections assessed by immunofluorescence (n = 4 mice per group, two slices per mouse). Scale bar 100 µm. **i,**Tight-junction proteins zonulin-1 (ZO-1) and occludin protein expression in homogenized brain tissue assessed by immunoblotting and **j**, quantification (in optical density; OD) (n = 6 per group). **k,** Brain microvascular structure of 3D vascular networks unveiled by brain clearing and **l**, quantification of fractional vascular volume and **m**, vascular length in Ctrl and GLUT1^1′GFAP^ mice (n = 3 mice per group). Data are presented as mean ± SEM. No significance was found as determined by two-tailed Student’s t test.

**Fig. S4.**
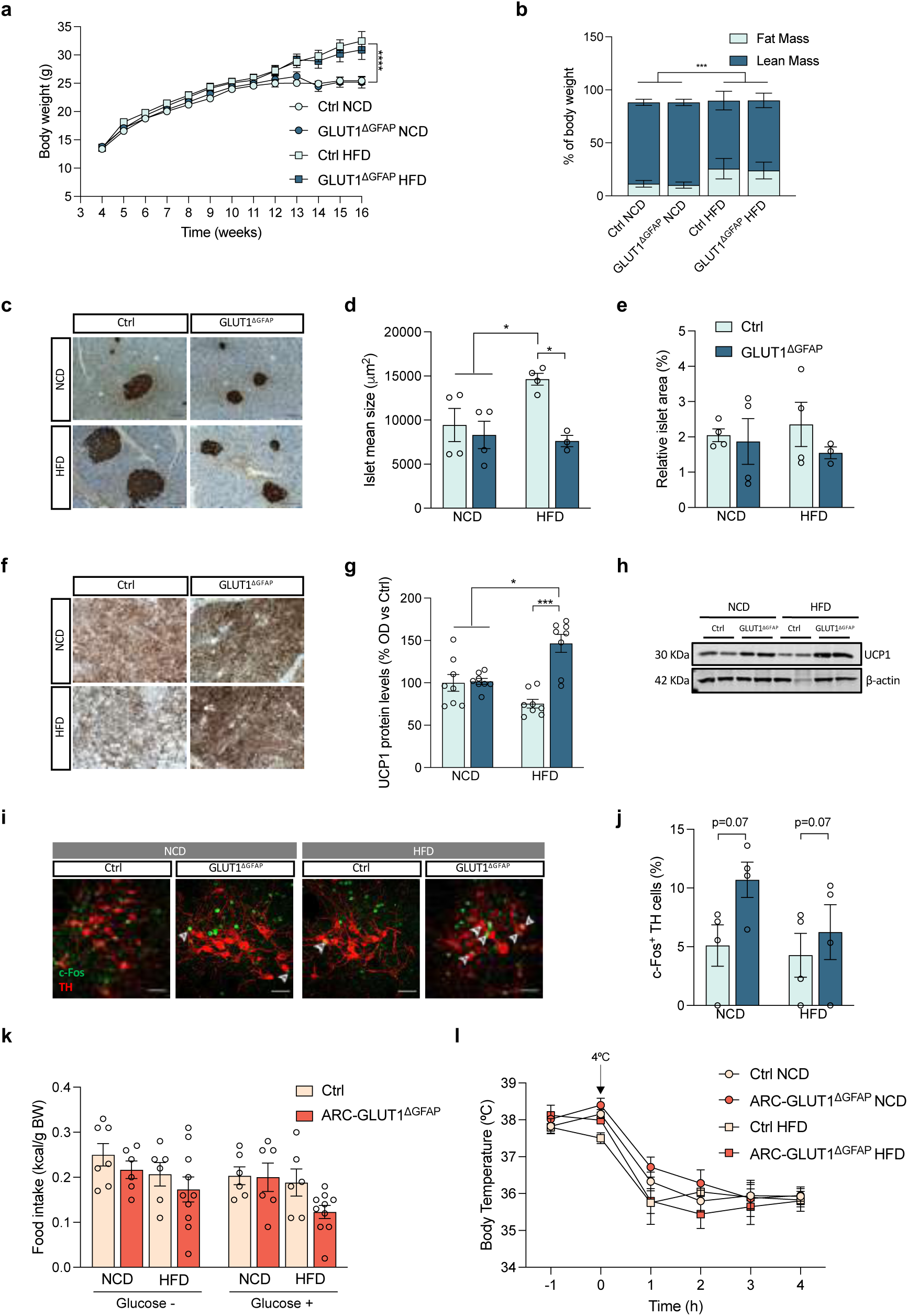
Astrocytic GLUT1 ablation induces peripheral glucose homeostasis improvement. **a**, weight and **b**, body composition assessment in normal chow diet (NCD)- and high fat diet (HFD)-fed control (Ctrl) and GLUT1^1′GFAP^ mice (n = 25-35 per group). **c,** Representative pancreatic islet image and **d**, quantification of islet mean size and **e**, relative islet area in control and GLUT1^1′GFAP^ mice in NCD or HFD (n = 4 mice per group, 10 islets per mouse). Scale bar 100 µm. **f,** Representative UCP1 protein staining, **g**, quantification of protein levels assessed by immunoblotting and **h**, a representative blot in normal chow diet (NCD)- and high fat diet (HFD)-fed control (Ctrl) and GLUT1^1′GFAP^ mice (n = 8 per group). Scale bar 100 µm. **i,** Representative c-Fos (green) staining of tyrosin hydroxylase (TH) positive (red) neurons of the locus coeruleus (LC) and **j**, quantification of c-Fos^+^-TH cells in control and GLUT1^1′GFAP^ mice in NCD or HFD (n = 4 mice per group, 2 slices per mouse). Scale bar 50 µm. **k**, Glucose-induced suppression of feeding evaluated by 4 h accumulated food intake in NCD- and HFD-fed Ctrl and ARC-specific astrocytic GLUT1 ablated (ARC-GLUT1^1′GFAP^) mice (n = 6-10 mice per group). **l,** Rectal temperature of control and ARC-GLUT1^1′GFAP^ mice on NCD and HFD upon cold exposure (+4°C) for 4 h (n = 7-10 mice per group). Data are presented as mean ± SEM. *p ≤ 0.05, ***p ≤ 0.001 as determined by repeated measurement ANOVA (a and l) and two-way ANOVA followed by Tukey (b, d, e, g, j, k).

**Fig. S5.**
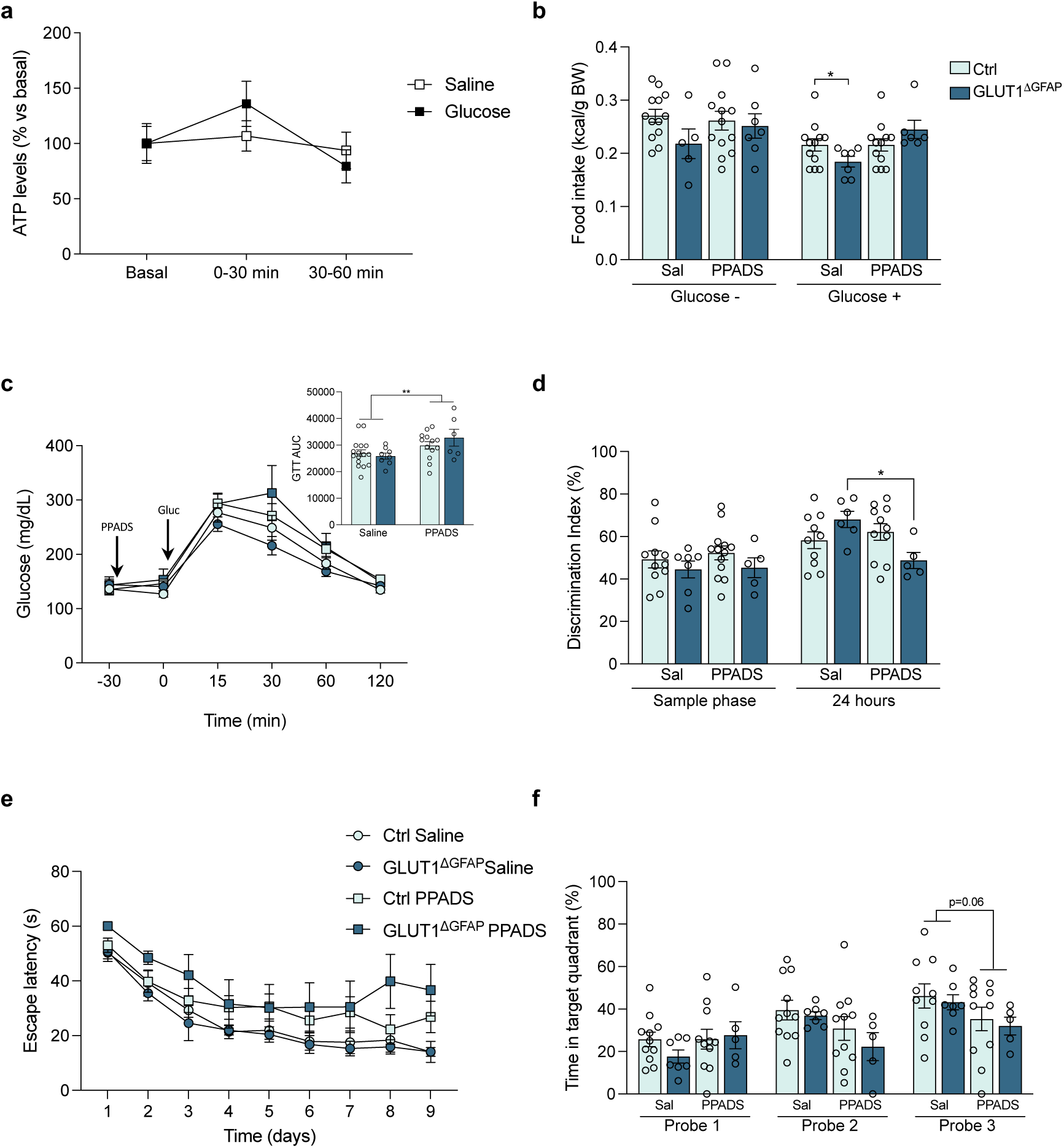
Purinergic signaling blocking abrogates the metabolic and cognitive benefits observed in GLUT1^1′GFAP^ mice. **a**, ATP levels in the microdialysis-isolated interstitial fluid of mice before (basal) and after i.p. saline or glucose injection (0-30 and 30-60 min). Data expressed as % of their own basal (n = 6 mice per group). **b**, Glucose-induced suppression of feeding evaluated by 4 h accumulated food intake in Ctrl and GLUT1^1′GFAP^ mice subjected to NCD after saline or PPADS i.c.v. injection (n = 6-13 per group). **c**, Glucose tolerance test in saline or PPADS i.c.v. treated Ctrl and GLUT1^1′GFAP^ mice on NCD and AUC quantification (n = 6-12 per group). **d**, Recognition memory assessment by novel object recognition (NOR) task of NCD-fed Ctrl and GLUT1^1′GFAP^ mice after saline or PPADS i.c.v. administration (n = 5-12 per group). **e**, Spatial memory evaluation by Morris water maze (MWM) acquisition phase and **f**, retention phase of saline or PPADS i.c.v. treated Ctrl and GLUT1^1′GFAP^ mice on NCD (n = 5-11 per group). Data are presented as mean ± SEM. *p ≤ 0.05 as determined by repeated measurement ANOVA (a, c and e) and two-way ANOVA followed by Tukey (b, c AUC, d and f).

**Fig. S6.**
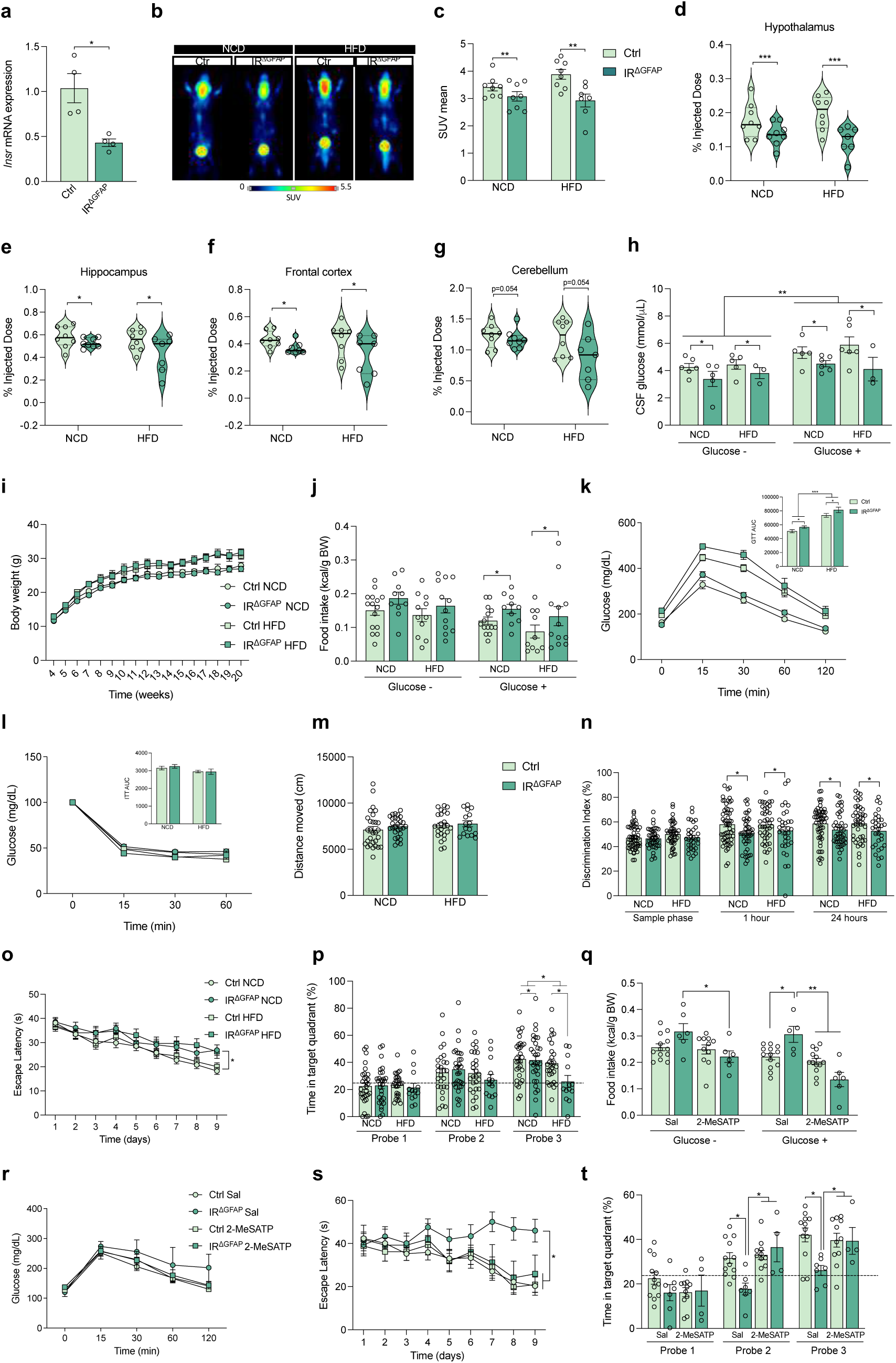
Astrocytic insulin receptor ablation induces metabolic and cognitive alterations that are rescued by purinergic signaling enhancement. **a**, *Insr* (insulin receptor coding gene) mRNA expression levels in control (Ctrl) and IR^1′GFAP^ mice (n = 4 mice per group). b, PET images of representative mice showing brain ^18^F-FDG signal, c, the quantification of positron emission (mean SUV) and d, *ex vivo* counting of radioactivity in dissected brain samples of hypothalamus, e, hippocampus, f, frontal cortex and g, cerebellum of Ctrl and IR^1′GFAP^ mice under normal chow diet (NCD) or high fat diet (HFD) (n = 8 mice per group). h, CSF glucose presence 30 min after i.p. vehicle or glucose injection in NCD- or HFD-fed Ctrl and IR^1′GFAP^ mice (n = 3-6 mice per group). i, Body weight assessment in NCD- and HFD-fed Ctrl and IR^1′GFAP^ mice (n = 25-35 per group). j, Accumulated food intake (4 h) after i.p. injection with vehicle or glucose in control and IR^1′GFAP^ mice under NCD or HFD (n = 10-15 per group). k, Glucose tolerance test (GTT) and l, insulin tolerance test (ITT) with the respective quantification of the area under the curve (AUC) in NCD- and HFD-fed Ctrl and IR^1′GFAP^ mice (n = 17-25 mice per group). m, Locomotor activity assessment in NCD- and HFD-fed Ctrl and IR^1′GFAP^ mice (n = 17-30 per group). n, Recognition memory assessment by novel object recognition (NOR) task performance of Ctrl and IR^1′GFAP^ mice. Data shows discrimination index (n = 32-40 per group). Spatial memory evaluation by Morris water maze o, acquisition phase and p, retention phase of Ctrl and GLUT1^1′GFAP^ mice in NCD and HFD (n = 20-40 per group). q, Glucose-induced suppression of feeding evaluated by 4 h accumulated food intake in Ctrl and IR^1′GFAP^ mice subjected to NCD after saline or 2-MeSATP i.c.v. injection (n = 6-13 per group). r, Glucose tolerance test in saline or 2-MeSATP i.c.v. treated Ctrl and IR^1′GFAP^ mice on NCD (n = 6-12 per group). s, Spatial memory evaluation by Morris water maze acquisition phase and t, retention phase of saline or 2-MeSATP i.c.v. treated Ctrl and IR^1′GFAP^ mice on NCD (n = 4-12 per group). Data are presented as mean ± SEM. *p ≤ 0.05, **p ≤ 0.01, ***p ≤ 0.001 as determined by two-tailed Student’s t test (a), two-way ANOVA followed by Tukey (c, d, e, f, g, h, j, k AUC, l AUC, m, n, p, q and t) and repeated measurement ANOVA (i, k, l, o, r and s).

**Fig. S7.**
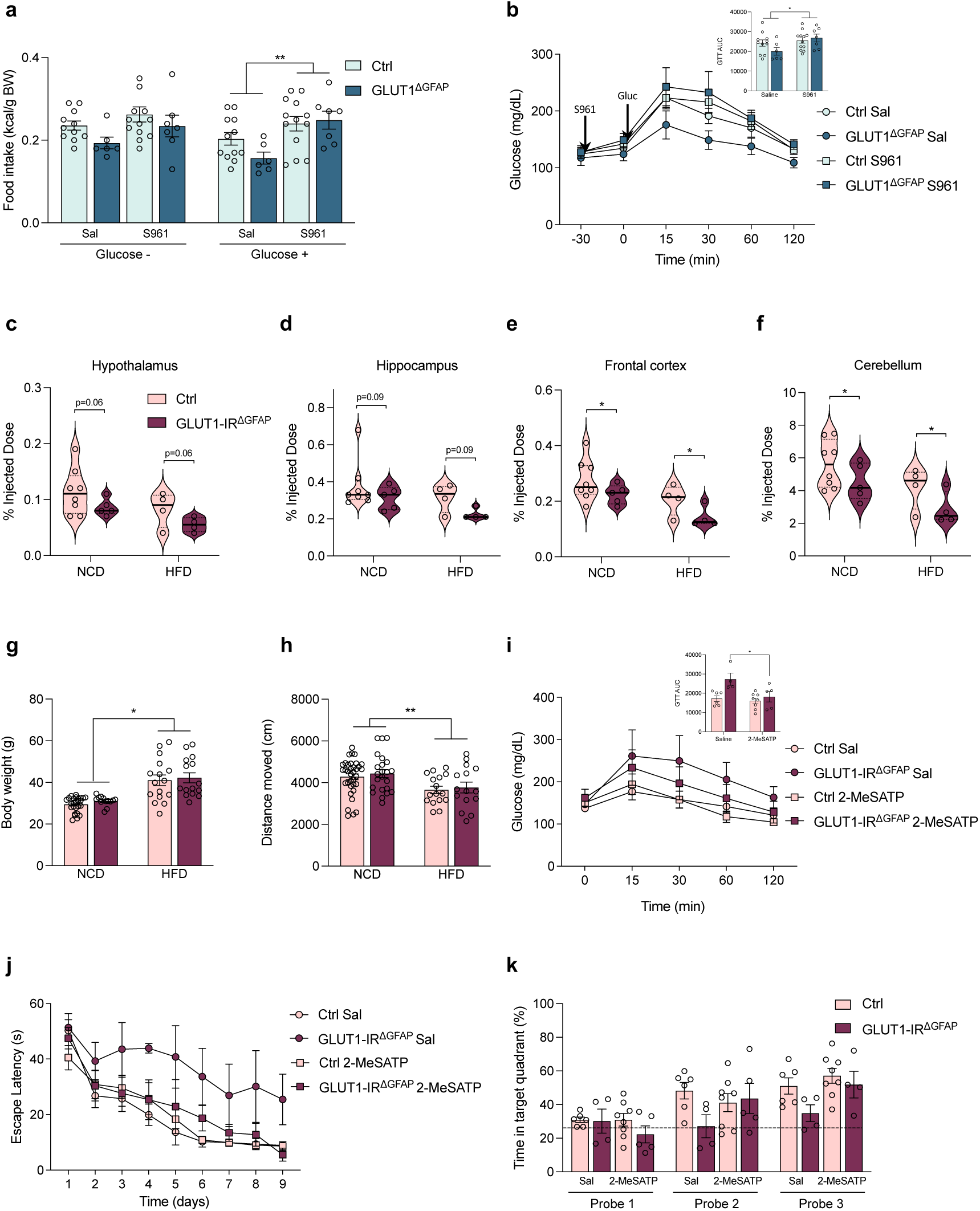
GLUT1-IR^1′GFAP^ mice showed metabolic and cognitive alterations are rescued by purinergic signaling enhancement. **a**, Glucose-induced suppression of feeding evaluated by 4 h accumulated food intake in Ctrl and GLUT1^1′GFAP^ mice subjected to NCD after saline or S961 i.c.v. injection (n = 6-12 per group). **b**, Glucose tolerance test in saline or S961 i.c.v. treated control and GLUT1^1′GFAP^ mice on NCD and the corresponding AUC quantification (n = 6-10 per group). **c**, *Ex vivo* counting of radioactivity in dissected brain samples of hypothalamus, **d**, hippocampus, **e**, frontal cortex and **f**, cerebellum of control (Ctrl) and GLUT1-IR^1′GFAP^ mice under normal chow diet (NCD) or high fat diet (HFD) (n = 4-8 mice per group). **g**, Body weight assessment in NCD- and HFD-fed Ctrl and GLUT1-IR^1′GFAP^ mice (n = 15-25 per group). **h,** Locomotor activity assessment in NCD- and HFD-fed Ctrl and GLUT1-IR^1′GFAP^ mice (n = 15-35 per group). **i,** Glucose tolerance test in saline or 2-MeSATP i.c.v. treated control and GLUT1-IR^1′GFAP^ mice on NCD (n = 5-8 per group). **j**, Spatial memory evaluation by MWM acquisition and **k**, retention phase of saline or 2-MeSATP i.c.v. treated control and GLUT1-IR^1′GFAP^ mice on NCD (n = 4-8 per group). Data are presented as mean ± SEM. *p ≤ 0.05, **p ≤ 0.01 as determined by two-way ANOVA followed by Tukey (a, b AUC, c, d, e, f, g, h, i AUC and k) and repeated measurement ANOVA (b, i and j).

**Supplementary Table 1.**
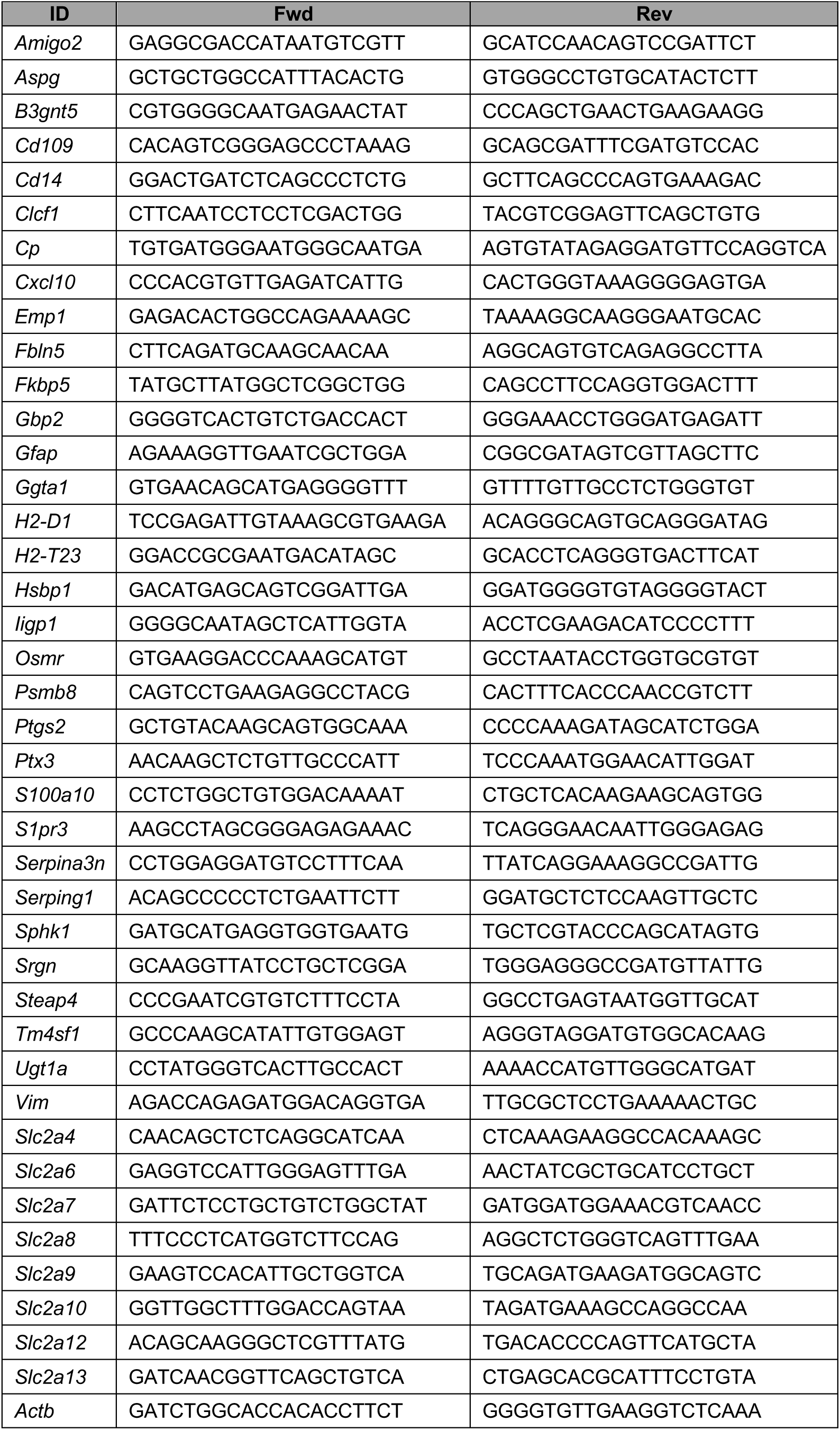
Primer sequences

